# A selective inference approach for FDR control using multi-omics covariates yields insights into disease risk

**DOI:** 10.1101/806471

**Authors:** Ronald Yurko, Max G’Sell, Kathryn Roeder, Bernie Devlin

## Abstract

To correct for a large number of hypothesis tests, most researchers rely on simple multiple testing corrections. Yet, new methodologies of selective inference could potentially improve power while retaining statistical guarantees, especially those that enable exploration of test statistics using auxiliary information (covariates) to weight hypothesis tests for association. We explore one such method, adaptive p-value thresholding (Lei & Fithian 2018, AdaPT), in the framework of genome-wide association studies (GWAS) and gene expression/coexpression studies, with particular emphasis on schizophrenia (SCZ). Selected SCZ GWAS association p-values play the role of the primary data for AdaPT; SNPs are selected because they are gene expression quantitative trait loci (eQTLs). This natural pairing of SNPs and genes allow us to map the following covariate values to these pairs: GWAS statistics from genetically-correlated bipolar disorder, the effect size of SNP genotypes on gene expression, and gene-gene coexpression, captured by subnetwork (module) membership. In all 24 covariates per SNP/gene pair were included in the AdaPT analysis using flexible gradient boosted trees. We demonstrate a substantial increase in power to detect SCZ associations using gene expression information from the developing human prefontal cortex (Werling et al. 2019). We interpret these results in light of recent theories about the polygenic nature of SCZ. Importantly, our entire process for identifying enrichment and creating features with independent complementary data sources can be implemented in many different high-throughput settings to ultimately improve power.

---

Large scale experiments, such as scanning the human genome for variation affecting a phenotype, typically result in a plethora of hypothesis tests. To overcome the multiple testing challenge, one needs corrections to limit false positives while maximizing power. Introduced by Benjamini & Hochberg (1995), false discovery rate (FDR) control has become a popular approach to improve power for detecting weak effects by limiting the expected false discovery proportion (FDP) instead of the more classical Family-Wise Error Rate. The Benjamini-Hochberg (BH) procedure was the first method to control FDR at target level *α* using a step-up procedure that is *adaptive* to the set of p-values for the hypotheses of interest (Benjamini & Hochberg 1995). Other methods for FDR control have led to improvements in power over BH by incorporating prior information, such as by the use of p-value weights (Genovese et al. 2006). In the “omics” world – genomics, epigenomics, proteomics, and so on – the challenge of multiple testing is burgeoning, in part because our ability to characterize omics features grows continually and in part because of the realization that multiple omics are required for describing phenotypic variation. One might imagine merging complementary omics data and tests using a priori hypothesis weights to improve power; however, until recently, it was not clear how to choose these weights in a data driven manner.

Recent methodologies have been proposed to account for covariates or auxiliary information while maintaining FDR control (Scott et al. 2015, Ignatiadis et al. 2016, Boca & Leek 2018, Li & Barber 2019, Zhang et al. 2019). We implement a selective inference approach, called adaptive p-value thresholding (Lei & Fithian 2018, AdaPT), to explore prior auxiliary information while maintaining guaranteed finite-sample FDR control. A recent review compared the performance of AdaPT with other covariate-informed methods for FDR control with off-the-shelf one and two-dimensional covariate examples (Korthauer et al. 2019). One of the weaknesses they ascribe to AdaPT is the unintuitive modeling framework for incorporating covariates; however, AdaPT is not a specific algorithm that one can simply apply to a dataset, but rather a meta-algorithm for marrying machine learning methods to multiple testing problems without compromising FDR control. We fully embrace AdaPT’s flexibility via gradient boosted trees in a much richer, high-dimensional setting. Our boosting implementation of AdaPT easily scales with more covariates, enabling practitioners to capture interactions and non-linear effects from the rich resources of prior information available.

In this manuscript, we demonstrate our gradient boosted trees implementation of AdaPT on results from genome-wide association studies (GWAS), incorporating covariates constructed from independent GWAS and gene expression studies. Specifically, we apply AdaPT to GWAS for detecting single nucleotide polymorphisms (SNPs) associated with schizophrenia (SCZ) using bipolar disorder (BD) GWAS results from an independent dataset as a covariate. Additionally, we incorporate results from the recent BrainVar study to identify a set of expression-SNPs (eSNPs) based on 176 neurotypical brains, sampled from pre- and post-natal tissue from the human dorsolateral prefrontal cortex (Werling et al. 2019). Along with the genetically correlated BD z-statistics, we create additional features from this complementary data source by summarizing the associated developmental gene expression quantitative trait loci (eQTL) slopes and membership in gene co-expression networks. We demonstrate that this process of identifying an enriched set of eSNPs and applying AdaPT with covariates summarizing gene expression from the developing human prefrontal cortex yield substantial improvement in power with each additional piece of information from the BrainVar study. Furthermore, we validate the replication of our results using more recent, independent SCZ studies.

This study had two goals, to explore the use of AdaPT in a realistic high-dimensional multiomics setting and to determine what can be learned about the neurobiology of SCZ by this exploration. Our results revealed the power of incorporating auxiliary information with flexible gradient boosted trees. While each covariate independently provided at best a modest increase in power, our adaptive search discovered a more complex model with far greater power. These discoveries also led to increasing support for the polygenic basis of SCZ, complementing recent findings and suggesting that there are many physiological avenues to its underlying neurobiology. We emphasize that the process and analysis undertaken with this implementation of AdaPT can be extended to a variety of “omics” and other settings to utilize the rich contextual information that is often ignored by standard multiple testing corrections. We highlight this feature by analyzing two other sets of GWAS studies, type 2 diabetes (T2D) and body mass index (BMI), using results from these analyses to interpret findings from SCZ.

## Results

### Methodology overview

AdaPT is an iterative search procedure, introduced by Lei & Fithian (2018), for determining a set of discoveries/rejections, 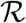, with guaranteed finite-sample FDR control at target level *α* under conditions outlined below. We apply AdaPT to the collection of p-values and auxiliary information, (*p*_*i*_, *x*_*i*_)_*i*∈*n*_, testing hypothesis *H_i_* regarding SNP *i*’s association with the phenotype of interest (e.g. SCZ). The covariates from some feature space, 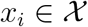, capture information collected independently of *p_i_*, but potentially related to whether or not the null hypothesis for *H*_*i*_ is true and the effect size under the alternative. AdaPT provides a flexible framework to incrementally *learn* these relationships, potentially increasing the power of the testing procedure, while maintaining valid FDR control.

For each step *t* = 0, 1, … in the AdaPT search, we first determine the rejection set 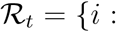 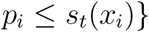, where *s_t_*(*x*_*i*_) is the rejection threshold at step *t* that is *adaptive* to the covariates *x*_*i*_. This provides us with both the number of discoveries/rejections 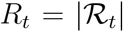, as well as a *pseudo*-estimate for the number of false discoveries *A*_*t*_ = |{*i* : *p*_*i*_ ≥ 1 − *s*_*t*_(*x*_*i*_)}| (i.e. number of p-values above the “mirror estimator” of *s*_*t*_(*x*_*i*_)). These quantities are used to estimate the FDP at the current step *t*,

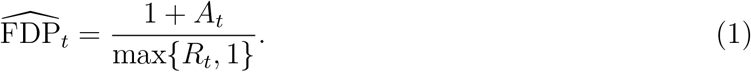

If 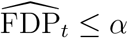, then the AdaPT search ends and the set of discoveries 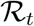 is returned. Otherwise, we proceed to update the rejection threshold while satisfying two protocols: (1) the updated threshold must be more stringent *s*_*t*+1_(*x*_*i*_) ≤ *s*_*t*_(*x*_*i*_), and (2) p-values determining *R*_*t*_ and *A*_*t*_ are *partially* masked,

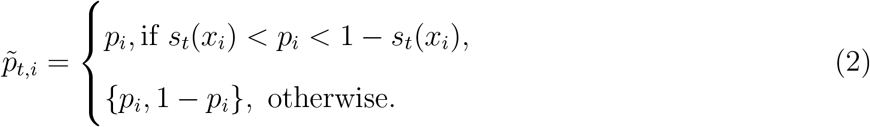

Under these protocols, the rejection threshold can be updated using *R*_*t*_, *A*_*t*_, and 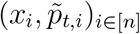. The flexibility in how this update takes place is one of AdaPT’s key strengths and allows it to easily incorporate other approaches from the multiple testing literature, such as a conditional version of the two-groups model (Efron et al. 2001) with estimates for the probability of being non-null, *π*_1_, and the effect size under the alternative, *µ*.

The algorithm proceeds by sequentially updating the threshold *s*_*t*+1_(*x*_*i*_) to discard the most likely null element in the current rejection region, as measured by the conditional local false discovery rate (fdr): i.e., 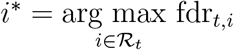 is removed from 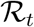. With the threshold updated, the AdaPT search repeats by estimating FDP and updating the rejection threshold until the target FDR level is reached 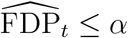 or 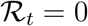.

This procedure guarantees finite-sample FDR control under independence of the null p-values and as long as the null distribution of p-values is *mirror conservative*, i.e. the large “mirror” counterparts 1 − *p*_*i*_ ≥ 0.5 are at least as likely as the small p-values *p*_*i*_ ≤ 0.5. To address the assumption of independence, we select a subset of weakly correlated SNPs detailed in *Data*, and additionally provide simulations in *SI Appendix* showing that AdaPT appears to maintain FDR control in relevant positive dependence settings. However, one practical limitation we encounter with the FDP estimate in Equation 1 is observing p-values *exactly* equal to one. While this can understandably occur with publicly available GWAS summary statistics, p-values equal to one will *always* contribute to the estimated number of false discoveries *A*_*t*_. This nuance can lead to a failure of obtaining discoveries at a desired target *α*, such as the reported AdaPT results by Korthauer et al. (2019) for multiple case-studies. However, we demonstrate in *SI Appendix* an adjustment to the p-values for T2D and BMI GWAS applications that alleviates this problem, although future work should explore modifications to the FDP estimator itself.

The modeling step of AdaPT estimates conditional local fdr with an EM algorithm. In this context, we use gradient boosted trees, which constructs a flexible predictive function as a weighted sum of many simple trees, fit using a gradient descent procedure that minimizes a specified objective function. The two objective functions considered correspond to estimating the probability of a test being non-null and the distribution of the effect size for non-null tests. The advantage of this approach to function fitting is that it is invariant to monotonic variable transformations, automatically incorporates important variable interactions, and is able to handle a large number of covariates without degrading significantly in performance due to the high dimensionality. In contrast, less effective methods might fail to capture useful information because the covariates are poorly transformed for a linear model, because the important information is only revealed through a combination of covariates, or because the important signal is simply swamped by the number of possible predictors to search through. Our choice of method gives the flexibility to include many potentially useful covariates without being overly concerned about the functional form with which they enter or their marginal utility. In our implementation, we employ the XGBoost library (Chen & Guestrin 2016) to capitalize on its computational advantages. Figure 1 displays the full pipeline of our implementation of AdaPT to GWAS summary statistics for SNPs using expression quantitative trait loci (eQTL) to select the SNPs under investigation.

**Figure 1:**
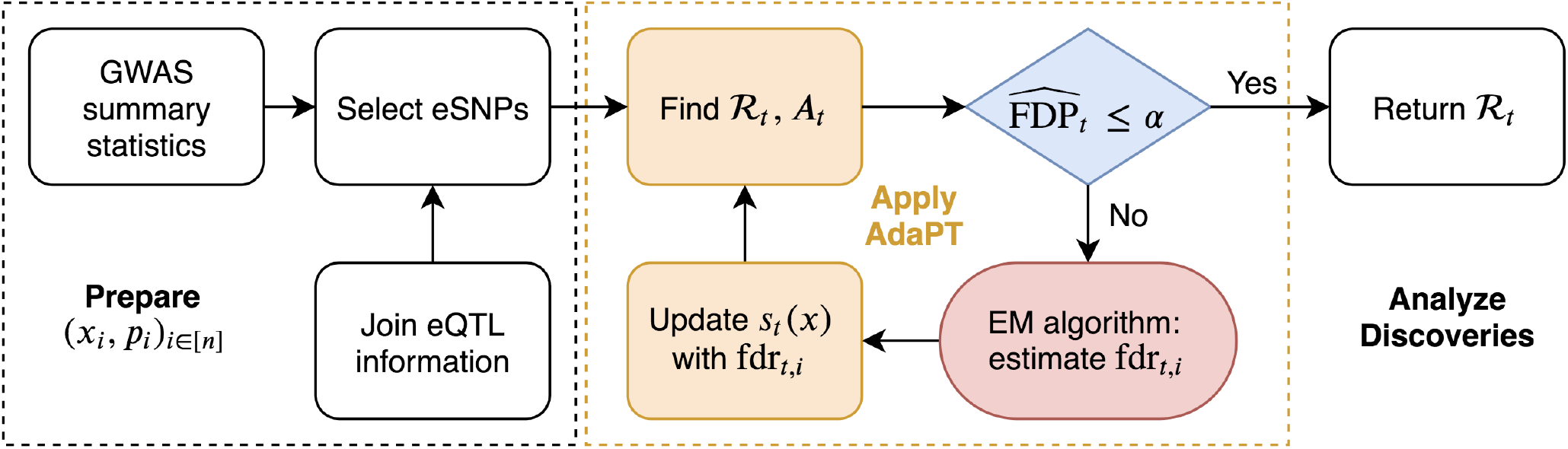
Summary of AdaPT GWAS implementation for selected set of SNPs. See Figure S1 for a summary of the AdaPT EM algorithm.

### Data

Our investigation includes AdaPT analyses of published GWAS p-values, {*p*_*i*_, *i* = 1, … *n*}, for body mass index (Locke et al. 2015, BMI), type 2 diabetes (Mahajan et al. 2018, T2D), and schizophrenia (Ruderfer et al. 2014, SCZ), but we focus our presentation on SCZ results. SCZ is a highly heritable, severe neuropsychiatric disorder. It is most strongly correlated, genetically, with another severe disorder, bipolar disorder (BD) (Lichtenstein et al. 2009, Cross-Disorder Group of the Psychiatric Genomics Consortium 2013). Because of this genetic correlation, reported z-statistics from BD GWAS, 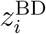, can be used as informative covariates for determining the SCZ rejection threshold. As an application of our AdaPT implementation, we use the GWAS summary statistics reported by Ruderfer et al. (2014), specifically 19,779 subjects diagnosed with either SCZ or BD with 19,423 control subjects (data are available from the Psychiatric Genomics Consortium, PGC). SCZ and BD subjects were completely independent and independent controls were bulk matched to the sample sizes of the two case samples. Results from more recent studies in Ruderfer et al. (2018) are used for replication analysis of our results (combined 53,555 SCZ and BD cases with 54,065 controls). However, the *2014-only* studies from Ruderfer et al. (2014) are a subset of the *all-2018* studies from Ruderfer et al. (2018). Although we do not have access to the raw genotype data, we use the fact that both papers report inverse variance-weighted fixed effects meta-analysis results (Willer et al. 2010). We then separate the summary statistics for the *2018-only* studies exclusive to Ruderfer et al. (2018), thus independent of the *2014-only* studies, and create an appropriate hold-out for replication analysis.

After matching alleles from both *2014-only* and *all 2018* studies and limiting SNPs to those with imputation score *INFO* > 0.6 for both BD and SCZ in *2014-only* (Ruderfer et al. 2014), we obtained 1,109,226 SNPs. Rather than test all SNPs, we chose to investigate a selected subset of SNPs, eSNPs, whose genotypes are correlated with gene expression; this additional filtering step captures a set of SNPs that are more likely to be functional and not highly correlated (Nicolae et al. 2010). These eSNPs were identified from two sources. First, we evaluated the BrainVar study of dorsolateral prefrontal cortex samples across a developmental span (Werling et al. 2019). BrainVar included cortical tissue from 176 individuals falling into two developmental periods: pre-natal, 112 individuals; and post-natal, 60 individuals. We identified *n*_SCZ_ = 25,076 eSNPs as any eQTL SNP-gene pairs provided by Werling et al. (2019) meeting Benjamini-Hochberg *α* ≤ 0.05 for at least one of the three sample sets (pre-natal, post-, and complete = all). These eSNPs were used for the SCZ analysis, which is a neurodevelopmental disorder and thus a developmental cohort seemed most appropriate for our analyses.

The second source was the Genotype-Tissue Expression (GTEx) V7 project dataset (GTEx Consortium 2015) with adult samples from fifty-three tissues. As the first winnowing step, we identified the set of GTEx eQTLs for *any* of the available tissues at target FDR level *α* = 0.05. Rather than use all GTEx eQTLs, however, we selected eQTL SNP-gene whose genotypes are most predictive of expression for each gene. The GTEx eSNPs were used for analysis of T2D and BMI, both of which typically onset in adults (for details see *SI Appendix*).

For each eSNP *i*, we created a vector of covariates *x*_*i*_ to incorporate auxiliary information collected independently of *p*_*i*_, including p-values from GWAS studies of related phenotypes, and relationships inferred from gene expression studies. First, we utilize the mapping of eSNPs to genes derived from eQTLs assessed in a relevant tissue type *r*. Although the majority of observed eSNPs have one unique cis-eQTL gene pairing, 14% of SNPs in BrainVar were eQTL for multiple genes. Let 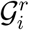 denote the set of genes whose expression is associated with eSNP *i* and summarize the level of expression as the average absolute eQTL slope for variants in 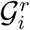 to obtain 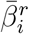. Additionally, we account for gene co-expression networks as covariates using the *J* = 20 modules reported in the BrainVar study, which were generated using weighted gene co-expression network analysis (Zhang & Horvath 2005, WGCNA). For each of the *j* = 1, … , *J* WGCNA modules, we create an indicator variable 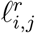 denoting whether or not eSNP *i* has *any* associated cis-eQTL genes in module *j*.

For the *n*_SCZ_ eSNPs, we calculate 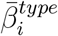 where *type* ∈ {pre, post, complete} to capture the eSNP’s overall expression association across different epochs of the developmental span. Additionally, we use the 20 WGCNA modules (including unassigned *gray*) reported by Werling et al. (2019) to create indicator variables 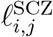 for *j* = 1, … , 20. This culminates in a vector of twenty-four covariates 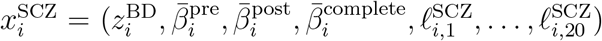. Although we use WGCNA modules to make use of the results from the BrainVar study, future applications could explore other approaches to account for gene set and pathway analysis (Zhu & Stephens 2018).

### AdaPT discoveries

As noted elsewhere (Schizophrenia Working Group of the Psychiatric Genomics Consortium 2014), eSNPs are more likely to be associated with a GWAS phenotype than are randomly chosen SNPs. This is true for the eSNP from BrainVar too, when evaluated in light of the SCZ GWAS p-values (Figure 2A). To evaluate the performance of the AdaPT search algorithm using the eSNP data, we compare the fitted full covariate model to results from its *intercept-only* version (Figure 2B versus 2C). As expected, the intercept-only analysis performs better than BH, with all 269 BH discoveries contained within the intercept-only discoveries, because it incorporates an estimate for the proportion of non-null tests. The full model rejects *R*_SCZ_ = 843 of the *n*_SCZ_ = 25,076 BrainVar eSNPs versus 361 discoveries for the intercept-only model. For insight into AdaPT’s performance, we sequentially include (1) only the BD z-statistics, then (2) include eQTL slope summaries, and then (3) the WGCNA indicators (Figure 2D-E). The largest number of discoveries occurs when all twenty-four covariates are fitted (Figure 2 D), highlighting that all three types of information *together* are required. Notably, only 540 associations are discovered using all covariates without interactions, fewer discoveries than only using module-based covariates with interactions. This highlights the improvement in AdaPT’s performance from modeling the interactions between covariates via gradient boosted trees. As might be expected from their counts of discoveries (Figure 2D), the greatest overlap with the full model occurs by fitting all covariates, but without interactions, or by fitting the module-based covariates (Figure 2E).

**Figure 2:**
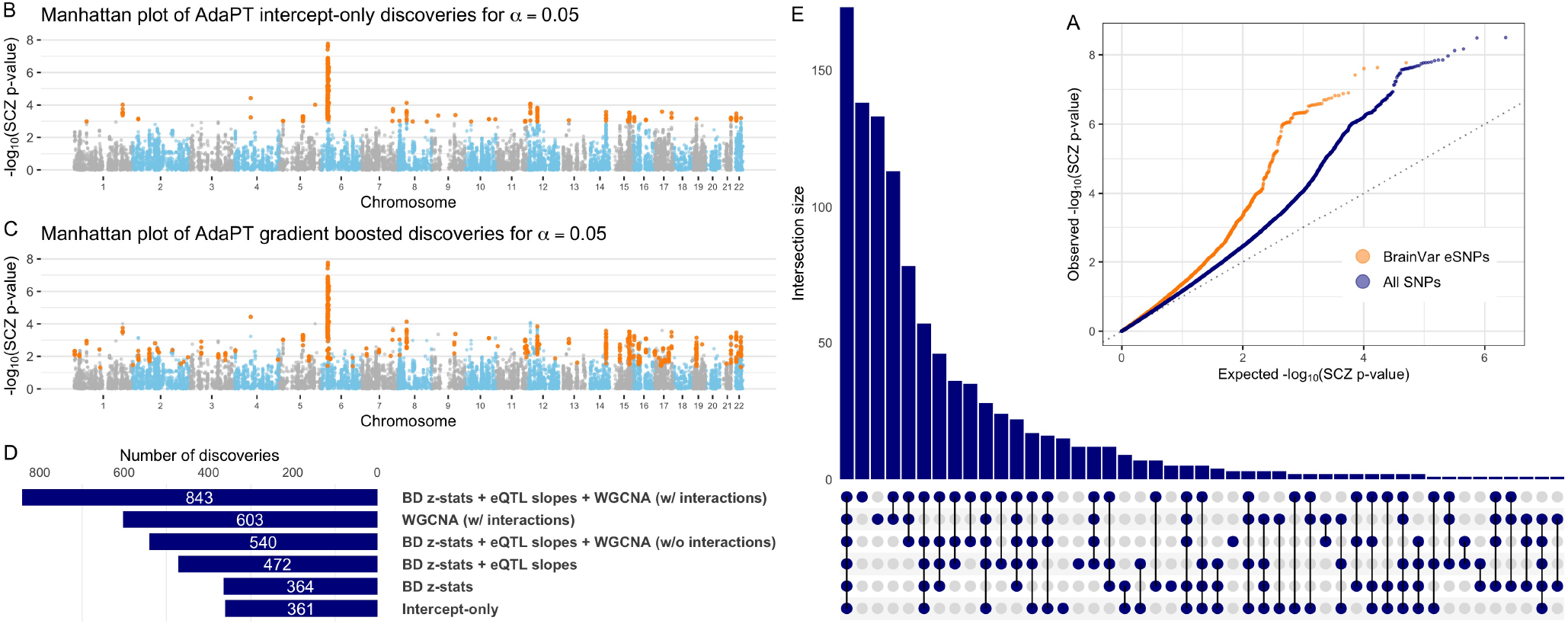
AdaPT results from analysis of schizophrenia (SCZ) p-values. *(A)* Comparison of qq-plots revealing SCZ enrichment for both BrainVar eSNPs compared to the full set of SNPs from *2014* studies. (B-C) Manhattan plots of SCZ AdaPT discoveries (in orange) using *(B)* intercept-only model compared to *(C)* covariate-informed model at target *α* = 0.05. *(D-E)* Comparison of the number of discoveries at target *α* = 0.05 for AdaPT with varying levels of covariates (D) and *(E)* their resulting discovery set intersections.

Additional discoveries are of little interest if they consist primarily of SNPs in LD with SNPs already discovered using a simpler model, such as the logit model typically used for SCZ GWAS. For context, however, of the initial 25,076 eSNPs we analyzed, only four have p-values < 5 × 10^−8^, the standard GWAS threshold, and all four occur in the discovery sets for the AdaPT full and intercept-only models. To investigate how the Adapt procedure performs using completely independent eSNPs, we identified the “lead” SNP in each LD block using the approach delineated in Schizophrenia Working Group of the Psychiatric Genomics Consortium (2014) and compared model performance for this set of approximately independent SNPs (*SI Appendix*). This thinning results in roughly 3,960 eSNPs to be analyzed by the different models (Figure 2). (Ties in q-values add or subtract a few SNPs to this 3,960 count, depending on the model analyzed.) When AdaPT is fit to these independent SNPs, we obtain analogous improvements in performance compared to the larger set of SNPs (Figures S2 and S3): the full AdaPT model discovers 95 independent loci, while the intercept-only model discovers only 42 loci. Likewise, the full model is the best model and interactions remain important. Finally, no location in the genome exerts unusual influence on the results, which is also the case for the analyses of 25,076 eSNPs.

As described previously, we performed similar analyses of T2D and BMI GWAS p-values. All results for these analyses, as well as more details regarding analyses of SCZ, are available in Dataset S1 and *SI Appendix*.

### Variable importance and relationships

We examine the variable importance and partial dependence plots from the gradient boosted models to provide insight into the relationships between each of the covariates and SCZ associations. Figure 3(A) displays the change in variable importance for the probability of being non-null (*π*_1_) at each model fitting iteration, with the top variables in the final model highlighted. We see that the BD z-statistics are estimated as the most important for each *π*_1_ model, but they decrease in importance in the final steps. In contrast, the unassigned *gray* module increases in important throughout the AdaPT search. This change in variable importance across the AdaPT search highlights that the difference in the discriminatory power of covariates depends on the remaining masked p-values.

**Figure 3:**
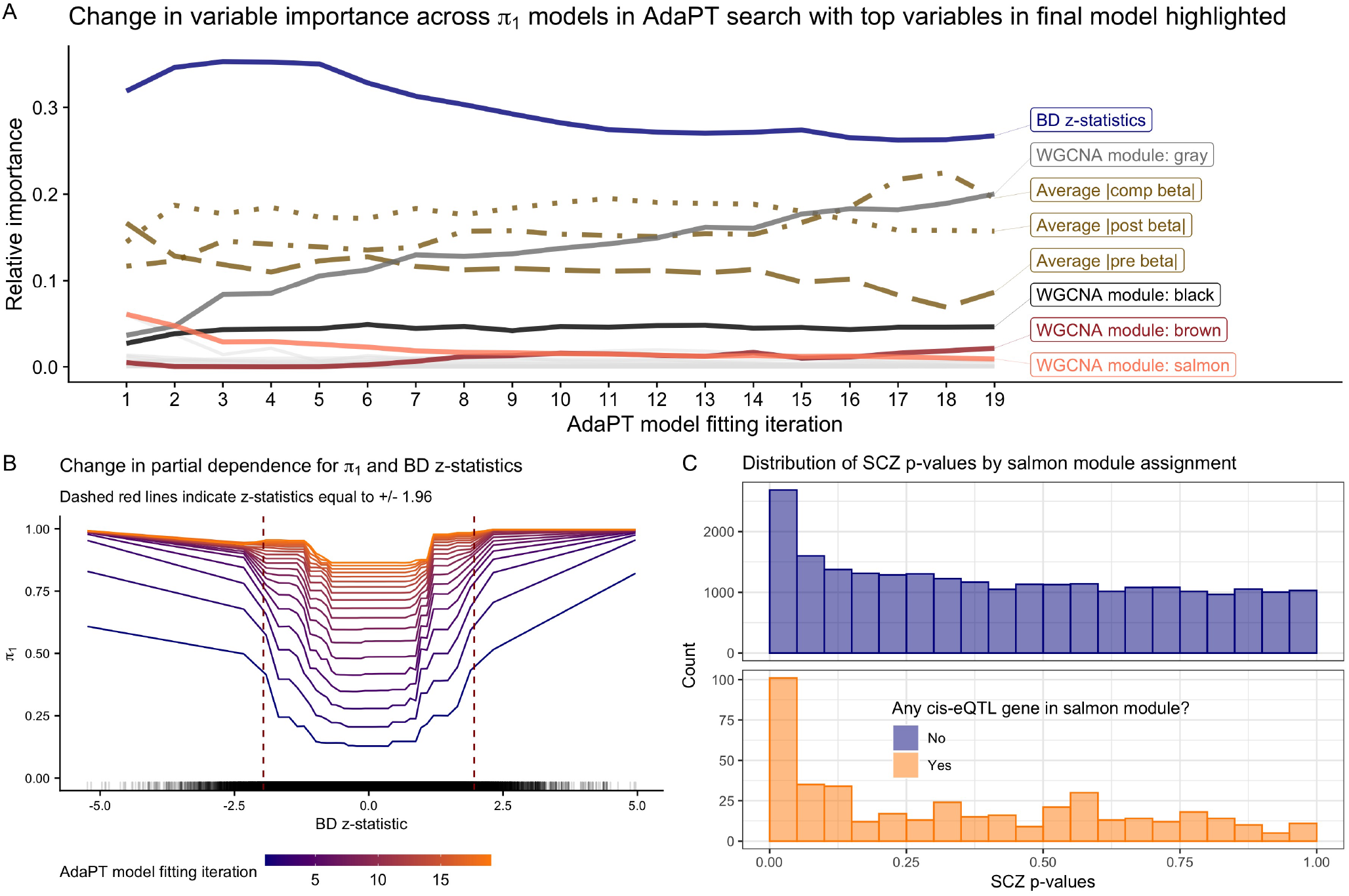
Variable importance and relationships. *(A)* Change in variable importance for AdaPT estimated probability of non-null *π*_1_ model across the search, with top variables in final model highlighted. *(B)* Change in partial dependence for estimated probability of being non-null *π*_1_ and BD z-statistics across *π*_1_ models in AdaPT search. *(C)* SCZ enrichment of eSNPs based on *salmon* WGCNA module membership, the most important WGCNA module indicator in the first model fitting step.

Figure 3(B) displays the partial-dependence plot (Friedman 2001) at each AdaPT model fitting iteration for the estimated marginal relationship between the BD z-statistics and the probability of being non-null, evaluated at the 0, 2.5%, 5%, … , 100% percentiles. Because the goal of the AdaPT two-groups model (detailed in *Methods*) is to order the remaining masked p-values, the *π*_1_ model predicts values relative to the remaining masked p-values: as the rejection threshold *s*_*t*_(*x*_*i*_) becomes more stringent, the masked p-values are more likely non-null (assuming there is signal). However, for each model iteration, Figure 3(B) reveals an increasing likelihood for non-null results as the BD z-statistics grow in magnitude from zero, as well as a diminished impact of BD z-statistics on the estimated *π*_1_ for later model iterations. Figure 3(C) displays the clear enrichment for eSNPs with cis-eQTL genes that are members of the *salmon* WGCNA module reported by Werling et al. (2019), which was the most important WGCNA module indicator in the first model fitting step. This differs from the unassigned *gray* module variable: it is predictive of SNPs that are classified as null, rather than associated with the phenotype. Taken together, Figures 3(A-C) emphasize the use of all covariates across different steps of the AdaPT search. See *SI Appendix* for more analyses highlighting the advantages of accounting for interactions between covariates.

### Replication in independent studies

Next, we examine the replicability of the *2014-only* SCZ AdaPT results using independent *2018-only* studies. We find (Figure 4) an increasing smoothing spline relationship between these sets of values, with noticeably increasing evidence indicated by the *2018-only* p-values for the set of AdaPT discoveries at *α* = 0.05. Additionally, of the 843 discoveries from the *2014-only* studies at target FDR level *α* = 0.05, approximately 55.2% (465 eSNPs) were nominal replications for *2018-only* (p-values < 0.05), comparable to the replication fraction expected on the basis of power (see *SI Appendix* for supporting simulations).

**Figure 4:**
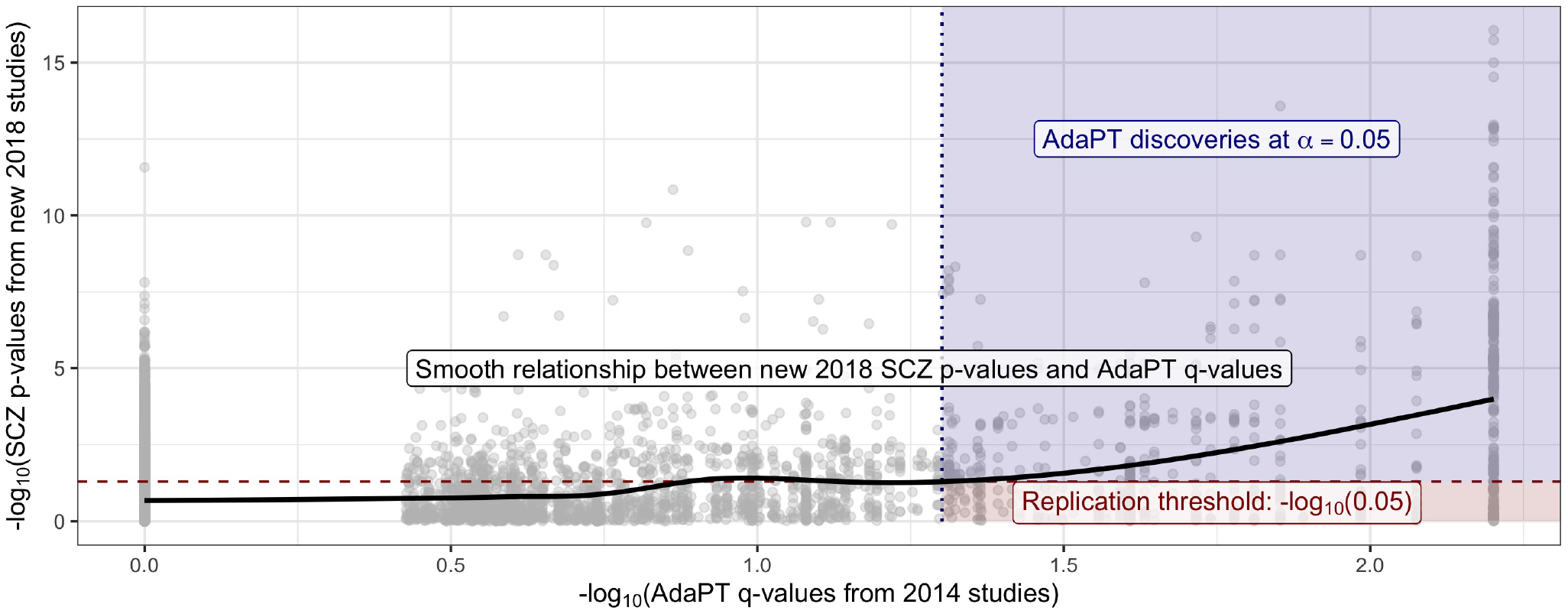
Relationship between the *2018-only* p-values and the resulting *2014-only* q-values from the AdaPT search. Black line displays smoothed relationship between SCZ p-values from *2018-only* studies and AdaPT q-values from *2014-only* studies. Blue region indicates AdaPT discoveries at *α* = 0.05 that are nominal replications, p-values from *2018-only* studies < .05, while red region denotes discoveries that failed to replicate.

### Gene ontology comparison

Using the SNP discoveries, which span the genome, we next sought biological insights. We applied gene ontology enrichment analysis (Ashburner et al. 2000, The Gene Ontology Consortium 2018) to the 136 genes obtained from the eQTL variant-gene pairs associated with the 843 discoveries. This analysis produced no clear signal, yielding only a minor enrichment for biological processes related to peptide antigen assembly. Several explanations are plausible, we explore two: either AdaPT is discovering SNPs of such small effect that the discoveries are not meaningful or SCZ is a highly complex disorder with a large number of biological processes involved. For comparison we applied our full pipeline to GWAS summary statistics for T2D (Mahajan et al. 2018). This comparison is of interest because T2D is a disease with a well understood functional basis and this is a well powered study (74,124 T2D cases and 824,006 controls). We restricted our analysis to 176,246 eSNPs based on eQTLs obtained using GTEx data. Next, we created eQTL-based covariates using pancreas, liver, and adipose tissue samples (see *SI Appendix* for more details). After creating a vector of covariates from GTEx, AdaPT returned 14,920 eSNPs at *α* = 0.05, resulting in 5,970 associated genes. Applying gene ontology enrichment analysis to this gene list, we discovered enrichment for biological processes related to lipid metabolic process (Figure 5), consistent with previous literature (Cirillo et al. 2018). These results provide some reassurance that the lack of specificity in the SCZ results can be attributed to the complex etiology of SCZ. For comparison to the well powered BMI GWAS (339,224 subjects), we found a lack of gene ontology enrichment in our gene discoveries (*SI Appendix*).

**Figure 5:**
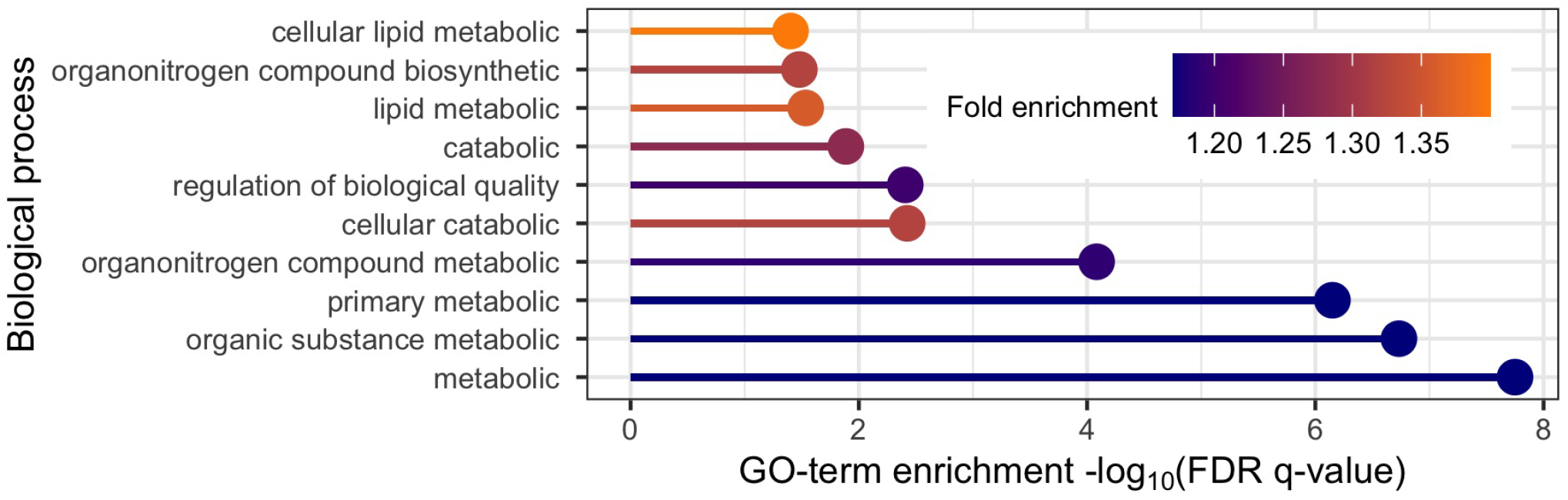
T2D gene ontology enrichment analysis results for top ten biological processes based on positive fold enrichment.

### Pipeline results for all 2018 studies

In addition to applying the pipeline to SCZ p-values from the *2014-only* studies in Ruderfer et al. (2014), we also modeled p-values from *all 2018* studies. The latter yields far more discoveries due to smaller standard errors from increased study sizes, even though the covariates were the same: for 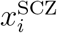, we find 2,228 discoveries at target FDR level *α* = 0.05 when the pipeline was applied to the p-values for most up-to-date set of studies versus 843 for the *2014-only* studies. Notably, the intercept-only version of AdaPT returned 1,865 discoveries at *α* = 0.05, meaning the covariates contributed to ≈ 19% increase in discovery rate for *all 2018* studies versus the ≈ 134% increase (361 to 843 eSNPs) from using the covariates for the *2014-only* studies. This reinforces the value of using auxiliary information in studies with lower power. Complementary to this observation, AdaPT applied to BMI GWAS the covariate informed models did not yield more discoveries than the intercept-only version (details presented in *SI Appendix*). Simply accounting for more auxiliary information does not guarantee an improvement in power and the advantages thereof diminishes as power increases, as witnessed by results for *all 2018* studies for SCZ and the large-scale BMI GWAS. Additionally, the larger number of discoveries for the SCZ *all 2018* studies, 2,228, maps onto 382 genes. Despite this increase, these genes did not reveal any clear signal from the Gene Ontology enrichment analysis, comporting with results from the *2014-only* results.

## Discussion

Our goals in this study were to explore the use of AdaPT for high-dimensional multi-omics settings and investigate the neurobiology of SCZ in the process. AdaPT was used to analyze a selected set of GWAS summary statistics for SNPs, together with numerous covariates. Specifically, SNPs were selected if they were documented to affect gene expression; these SNP-gene pairs were dubbed eSNPs. Covariates for these eSNPs included GWAS test statistics from a genetically correlated phenotype, BD, which were mapped to eSNPs through SNP identity; as well as features of gene expression and co-expression networks, which were mapped to eSNPs through genes. By coupling flexible gradient boosted trees with the AdaPT procedure, relationships among eSNP GWAS test statistics and covariates were uncovered and more SNPs were found to be associated with SCZ, while maintaining guaranteed finite-sample FDR control. The tree-based handling of covariates addresses a perceived weakness of AdaPT, namely the unintuitive modeling framework for incorporating covariates (Korthauer et al. 2019). Moreover, it is worth noting that the original approach implemented by Lei & Fithian (2018), a generalized linear model with spline bases, yields similar results (361 discoveries at target *α* = 0.05) when applied to the univariate case of only using BD z-statistics. This is an even more straightforward implementation for handling covariates without interactions. The pipeline we built should be simple to mimic for a wide variety of omics and other analyses.

The results shed light on the level of complexity underlying the neurobiology of SCZ. If the origins of SCZ arose by perturbations of one or a few pathways, we would expect to converge on those pathways as we accrue more and more genetic associations. On the other hand, if the ways to generate vulnerability to SCZ were myriad — even if there is an single ultimate cause shared across all cases — then we might expect no such convergence, at least with regards to the common variation assessed through GWAS. Gene ontology analysis of associated discovery genes from either the *2014-only* or *all 2018* studies reveals no enrichment for biological processes for SCZ. There are many possible explanations for these null findings, one of which is simply a lack of power or specificity of our results. However, the result stands in stark contrast to the results for T2D, for which the gene ontology analysis converges nicely on accepted pathways to T2D risk; yet they comport with those for BMI, which is known to have myriad genetic and environmental origins. Therefore our results are consistent with myriad pathways to vulnerability for SCZ, although it is impossible to rule out other explanations: for example, the possibility that we understand so little about brain functions that gene ontology analyses lack specificity. In any case, our results are consistent with two recent theories underlying the genetics of SCZ, namely extreme polygenicity (O’Connor et al. 2019) and “omnigenic” origins (Boyle et al. 2017).

Although the examples considered in this manuscript pertain to omics data, this process can be adapted for a large variety of settings. We demonstrate in *SI Appendix* simulations showing that AdaPT appears to maintain FDR control in positive dependence settings emulating linkage disequilibrium (LD) block structure underlying GWAS results. There is a clear need, however, for future work to explore AdaPT’s properties and computational challenges under various dependence regimes. The growing abundance of contextual information available in “omics” settings provides ample opportunity to improve power for detecting associations, using a flexible approach such as AdaPT, when addressing the multiple testing challenge.

## Methods

### Two-groups model

The most critical step in the AdaPT algorithm (Lei & Fithian 2018) involves updating the rejection threshold *s*_*t*_(*x*_*i*_). Following Lei & Fithian (2018), we use a conditional version of the classical two-groups model (Efron et al. 2001, Scott et al. 2015) where the null p-values are modeled as uniform (*f*_0_(*p*|*x*) ≡ 1) and we model the non-null p-value density with a beta distribution density parametrized by 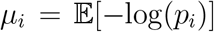, resulting in a conditional density for a beta mixture model, 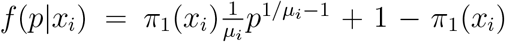. In this form, we can model the non-null probability 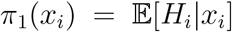 and the effect size for non-null hypotheses 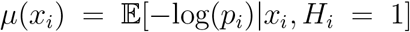 with two separate gradient boosted tree-based models. The XGBoost library (Chen & Guestrin 2016) provides logistic and Gamma regression implementations which we use for *π*_1_(*x*_*i*_) and *µ*(*x*_*i*_) respectively.

There are two categories of missing values in these regression problems: *H*_*i*_ is never observed, and at each step *t* of the search, the p-values for tests {*i* : *p*_*i*_ ≤ *s*_*t*_(*x*_*i*_) or *p*_*i*_ ≥ 1 − *s*_*t*_(*x*_*i*_)} are masked as 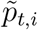. An expectation-maximization (EM) algorithm can be used to estimate both 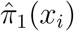 and 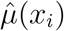 by maximizing the partially observed likelihood. We briefly restate the EM algorithm from (1), and provide details in our supplementary materials that reflect the approach taken in the `R adaptMT` package by the same authors, which differs slightly from (1).

During the E-step of the *d* = 0, 1, … iteration of the EM algorithm, conditional on the partially observed data fixed at step *t*, 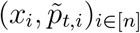, we compute both, 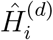 and 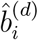, where 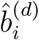 indicates how likely 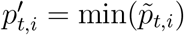 equals *p*_*i*_ for non-null hypotheses. The explicit calculations of 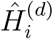 and 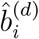 are available in the supplementary materials of Lei & Fithian (2018).

The M-step consists of estimating 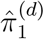 and 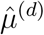 with separate gradient boosted trees, using

*pseudo*-datasets to handle the partially masked data. In order to fit the model for *π*_1_(*x*_*i*_), we construct the response vector 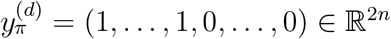 and use weights 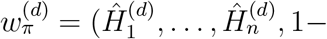 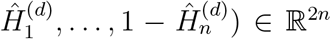. Then we estimate 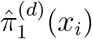 using the first *n* predictions from a classification model using 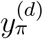 as the response variable with the covariate matrix (*x*_*i*_)_*i*∈[*n*]_ replicated twice and weights 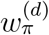. Similarly, for estimating 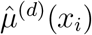 we construct a response vector 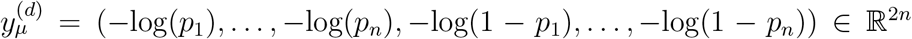 with weights 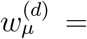 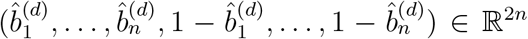, and again take the first *n* predicted values using the duplicated covariate matrix.

We follow the procedure detailed in Section 4.3 of Lei & Fithian (2018) to estimate the conditional local fdr for each 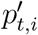, and then update the rejection threshold to *s*_*t*+1_(*x*_*i*_) by removing test 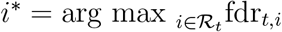 from 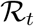.

### AdaPT gradient boosted trees with CV steps

As a flexible approach for modeling the conditional local fdr, we use gradient boosted trees (Friedman 2001) via the open-source XGBoost implementation (Chen & Guestrin 2016). Gradient boosted trees are an ensemble of many small tree models that jointly contribute to predictions.

Let 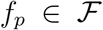 be an individual regression tree, then the sum-of-trees model can be written as, 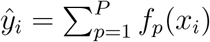 to minimize 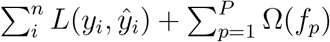 where *L* is the loss function and Ω measures the complexity of each tree such as the maximum depth, regularization, etc. Chen & Guestrin (2016) detail the algorithms for fitting the model in an additive manner as well as determining the splits for each tree.

To tune the variety of parameters for gradient boosted trees within AdaPT, such as the number of trees *P* and maximum depth of each tree, we use the cross-validation (CV) approach recommended in Lei & Fithian (2018). If we are considering *M* different options of boosting parameters, then we evaluate each of the *M* choices during the modeling phase of the AdaPT search. At step *t*, we divide the data into *K* folds preserving the relative proportions of masked and unmasked hypotheses. Then for each set of boosting parameters *m* = 1, …, *M*, and for each fold *k* = 1, … *K*: (1) apply EM-algorithm to estimate 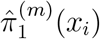 and 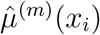 using parameters *m* with data from folds {1, … , *K*}\{*k*}, and (2) compute expected-loglikelihood 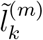 on hold-out set *k* using two-groups model parameters from *m* following convergence, and compute total across folds as 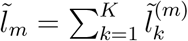. Finally we use the set of parameters 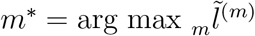 in another instance of the EM algorithm to estimate 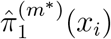 and 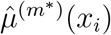 on all data.

### Computational aspects of AdaPT

Practical decisions are necessary to implement the AdaPT search. In addition to the covariates and p-values (*x*_*i*_, *p*_*t,i*_)_*i*∈[*n*]_, an initial rejection threshold *s*_0_(*x*_*i*_) is required to begin the search. Rather begin the search with a high starting threshold, such as 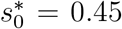 recommended by Lei & Fithian (2018), we instead begin the AdaPT search with 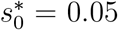. Our decision to lower the starting threshold is advantageous for multiple reasons. First, intuitively, this starts our search in the regime of interest for target level *α* = 0.05, whereas we would not expect to detect discoveries with larger p-values using this flexible multiple testing correction. Additionally, by lowering the starting threshold, more true information is available to the gradient boosted trees at the start of the AdaPT search. For instance, with the set of BrainVar eSNPs, 21,248 true p-values are immediately revealed with 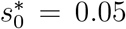 as compared to only 2,290 when 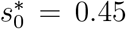. Simulations detailed in *SI Appendix* show that on average our choice for using a lower threshold results in higher power.

The most computationally intensive part of the procedure is updating the rejection threshold via the EM algorithm. Instead of updating the model for estimating fdr_*t,i*_ at each step of the search, we re-estimate every [*n*/20] steps as recommended by Lei & Fithian (2018). However, the inclusion of the previously described *K*-fold CV procedure (we use *K* = 5) for tuning the gradient boosted trees obviously adds computational complexity to the AdaPT search, and would be expensive to apply every time the model fitting takes place. Rather, we apply the CV step once at the beginning, and then another time half-way through the search based on the similarity of simulation performance with varying number of CV steps in *SI Appendix*. Additionally, one needs to choose the potential *M* model parameter choices. Technically, unique combinations can be used for both models, *π*_1_ and *µ*, but for simplicity we only consider matching settings for both models, i.e. both models have the same number of trees and maximum depth. As a reminder, AdaPT guarantees finite-sample FDR control **regardless** of potentially over-fitting to the data when using the CV procedure. Simulations are provided in *SI Appendix* showing how extensively increasing the number of trees *P* leads to decreasing power, but maintains valid FDR control.

We provide a modified version of the `adaptMT R` package to implement the AdaPT-CV tuning steps with XGBoost models at https://github.com/ryurko/adaptMT, and provide all code used to generate the manuscript’s results at: https://github.com/ryurko/AdaPT-GWAS-manuscript-code.

## Acknowledgements

This work was supported by National Institute of Mental Health Grant R37MH057881, the Simons Foundation Grant SFARI SF575097, and the National Science Foundation DMS 1613202 Grant. We are grateful for help from Lambertus Klei regarding data sets and their interpretation.

## Supporting Information Text

### AdaPT conditional two-groups model

This section provides a more detailed explanation of updating the rejection threshold *s*_*t*_(*x*_*i*_) in the AdaPT procedure, expanding on the description from *Methods* in the main manuscript. As in the main text, this is essentially an explanation of the EM approach of Lei & Fithian (2018). Note that for coherence some text is repeated from the main manuscript. Lei & Fithian (2018) use a conditional version of the classical two-groups model (Efron et al. 2001) yielding the conditional mixture density,

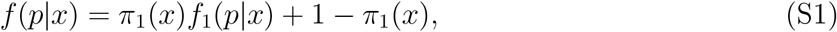

where the null p-values are modeled as uniform (*f*_0_(*p|x*) ≡ 1). They proceed to use a *conservative* estimate for the conditional local false discovery rate, 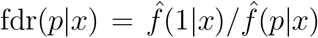, by setting 1 − *π*_1_(*x*) = *f* (1|*x*).

We model the non-null p-value density with a beta distribution density parametrized by *µ*_*i*_,

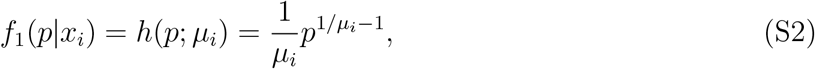

where 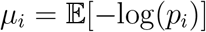, resulting in a conditional density for a beta mixture model,

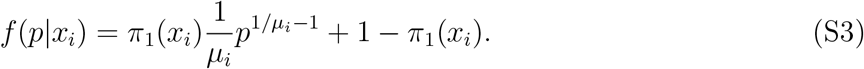

In this form, we can model the non-null probability 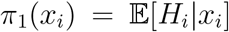 and the effect size for non-null hypotheses 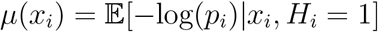 with two separate gradient boosted tree-based models. The XGBoost library (Chen & Guestrin 2016) provides logistic and Gamma regression implementations which we use for *π*_1_(*x*_*i*_) and *µ*(*x*_*i*_) respectively.

There are two categories of missing values in these regression problems: *H*_*i*_ is never observed, and at each step *t* of the search, the p-values for tests {*i* : *p*_*i*_ ≤ *s*_*t*_(*x*_*i*_) or *p*_*i*_ ≥ 1 − *s*_*t*_(*x*_*i*_)} are masked as 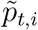. An expectation-maximization (EM) algorithm can be used to estimate both 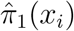 and 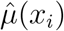 by maximizing the partially observed likelihood. The complete log-likelihood for the conditional two-groups model is,

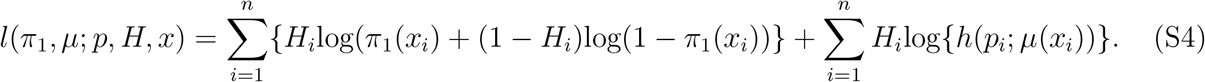

During the E-step of the *d* = 0, 1, … iteration of the EM algorithm, conditional on the partially observed data fixed at step *t*, 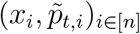, we compute both,

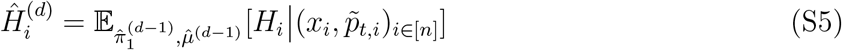

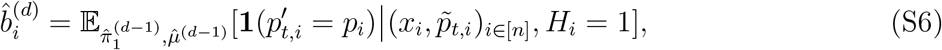

where 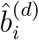 indicates how likely 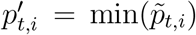 equals *p*_*i*_ for non-null hypotheses. The explicit calculations of 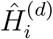 and 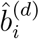 for both the revealed, 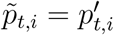, and masked p-values, 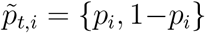, are available in the supplementary materials of Lei & Fithian (2018).

The M-step consists of estimating 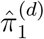 and 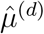 with separate gradient boosted trees, using *pseudo*-datasets to handle the partially masked data. In order to fit the model for *π*_1_(*x*_*i*_), we construct the response vector 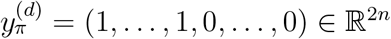 and use weights 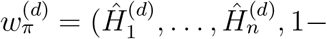 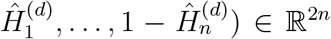. Then we estimate 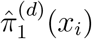 using the first *n* predictions from a classification model using 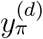 as the response variable with the covariate matrix (*x*_*i*_)_*i*∈[*n*]_ replicated twice and weights 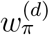. Similarly, for estimating 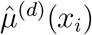 we construct a response vector 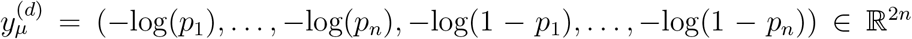 with weights 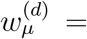 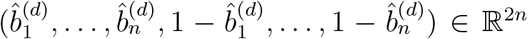, and again take the first *n* predicted values using the duplicated covariate matrix.

The conditional local fdr is estimated for each 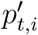,

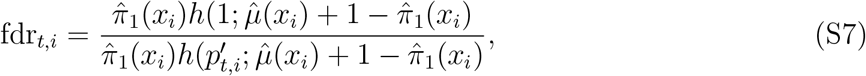

and we follow the procedure detailed in Section 4.3 of Lei & Fithian (2018) to update the rejection threshold to *s*_*t*+1_(*x*_*i*_) by removing test 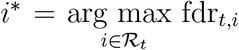 from 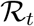. A summary diagram of the EM algorithm is displayed in Figure S1.

### SCZ results with independent loci

One potential concern regarding the assessment of performance of AdaPT is the impact of linkage disequilibrium (LD). In the Manhattan plots of Figures 2(B-C), the discoveries visually appear to be located close to one another. However, the visual appearance of genomic positions is somewhat misleading because our initial selection of eSNPs greatly reduces the number of SNPs commonly portrayed in Manhattan plots many of these SNPs are not very close to each other in the genome and not in high LD, although the format of the Manhattan plot makes this feature hard to see. To take this analysis further we follow common practice for GWAS results by identifying the “best” or “lead” SNPs in a LD block/cluster, using a similar approach as Schizophrenia Working Group of the Psychiatric Genomics Consortium (2014), for each of the set of discoveries presented in Figure 2:

1. order the SNPs by the AdaPT −log_10_(q-value) in descending order,
2. starting with the SNP with the largest value for the AdaPT −log_10_(q-value),
  - remove all SNPs with *r*^2^ ≥ 0.1 within a 500kb window,
  - move on to next SNP that is still remaining,
3. return the retained SNPs as the LD-independent SNPs in low LD (*r*^2^ < 0.1). (Remark: this approach excludes SNPs whose contribution to the GWAS signal is partially independent of the lead SNP, but it has the advantage of simplicity.)

**Figure S1:**
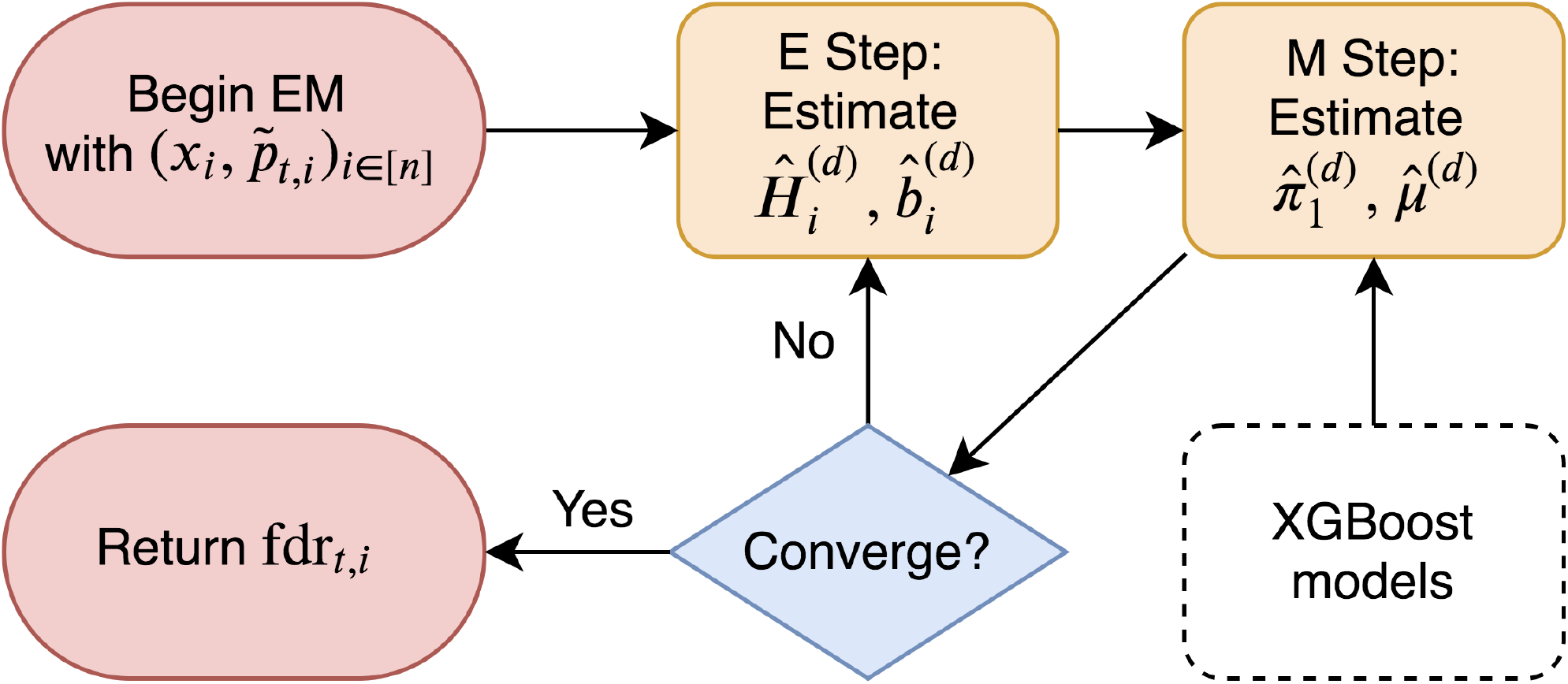
Summary of AdaPT EM algorithm.

We use the reference European sample genotype data from the 1000 Genomes project (1000 Genomes Project Consortium and others 2012) to compute the *r*^2^ values between SNPs. In the GWAS setting this LD clumping procedure is typically applied to the reported SNP p-values, but because the ordering of SNPs varies between the different sets of discoveries (intercept-only versus use of covariates) we perform the operation separately with their respective q-values. For each of the different set of covariates considered, this results in reducing the 25,076 selected eSNPs down to the following number of “independent loci”:

- Intercept-only: 3,958
- BD z-stats: 3,966
- BD z-stats + eQTL slopes: 3,962
- BD z-stats + eQTL slopes + WGCNA (w/ interactions): 3,963
- BD z-stats + eQTL slopes + WGCNA (w/o interactions): 3,959
- WGCNA: 3,954

The differences in counts are due to the different number of ties that take place between the resulting q-values for each considered set of covariates. Next, for the identified set of “lead” SNPs we observe how many have q-values less than the target FDR level *α* = 0.05 (i.e. associations detected at *α* = 0.05). The results are displayed in the Figure S2, including Manhattan plots Figures S2(A-B) of the q-values for the AdaPT intercept-only and BD z-stats + eQTL slopes + WGCNA (w/ interactions) results, rather than using the actual p-values. The lead SNPs in each of the Manhattan plots are denoted by an X shape. In conjunction with Figures S2(C-D), the relative improvement in the set of independent loci within the discovery sets from AdaPT is analogous to the results presented in Figure 2, emphasizing the advantage of accounting for covariates and their interactions via gradient boosted trees. Additionally, Figure S3 further emphasizes that the improvement in power is not restricted to a particular section of the genome. As seen in Figure S4, we observe a similar improvement in the number of independent loci when ordering the SNPs with the observed 2014-only studies SCZ p-values.

While we maintain FDR control on the original set of discoveries (see Figure 3 in *Results*), we do not retain any guarantees regarding the detected independent loci presented in Figure S2. In order to maintain FDR control on the set of discovered independent loci, an alternative approach or adjustment to the AdaPT algorithm is required. A simple alternative is to first apply LD pruning/clumping as initial step prior to applying AdaPT to a reduced set of lead SNPs. However, this encounters the challenge of defining lead SNPs without data “snooping” based on using the observed p-values. Future work will explore modifications for AdaPT, potentially exploring recent developments (Ren & Candès 2020), to maintain FDR control on an independent subset of SNPs.

**Figure S2:**
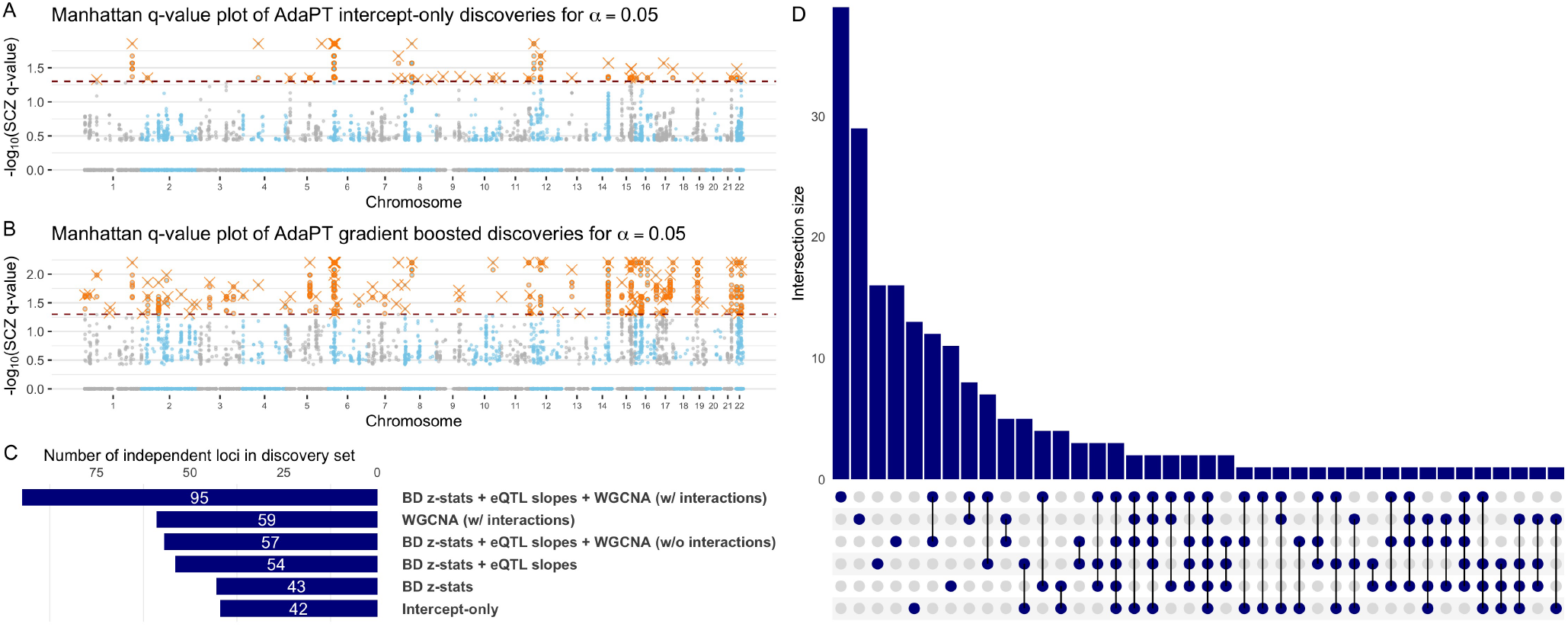
Manhattan q-value plots of SCZ AdaPT discoveries (orange) using *(A)* intercept-only model compared to *(B)* covariate informed model at target *α* = 0.05, with lead SNPs for independent loci denoted by Xs. *(C)* Comparison of the number of independent loci for each discovery set at target *α* = 0.05 based on LD pruning with the respective AdaPT q-values and *(D)* their resulting discovery set intersections.

### SCZ variable importance and partial dependence

We explore further the variable relationships from the gradient boosted trees. First, Figure S5 displays the change in variable importance for the non-null effect size (*µ*) at each model fitting iteration, with the top variables in the final model highlighted. The variable importance measures are relatively stable across all model iterations with the BD z-statistics and eQTL slope measures maintaining the highest level of importance. Figure S6 displays the partial-dependence plot at each AdaPT model fitting iteration for the estimated marginal relationship between the BD z-statistics and the non-null effect size *µ*, evaluated at the 0, 2.5%, 5%, … , 100% percentiles. The estimates reveal an increasing effect size as the BD z-statistics grow in magnitude, which is relatively stable across the model iterations. Figures S7(A-C) display the relationships for the probability of non-null model, while (D-F) display relationships for the effect size under the alternative. Although the partial dependence plots show considerable variability due to the high dimensional of the model, we can still see general trends consistent with the variable importance plots from Figure 3(A) and Figure S5.

**Figure S3:**
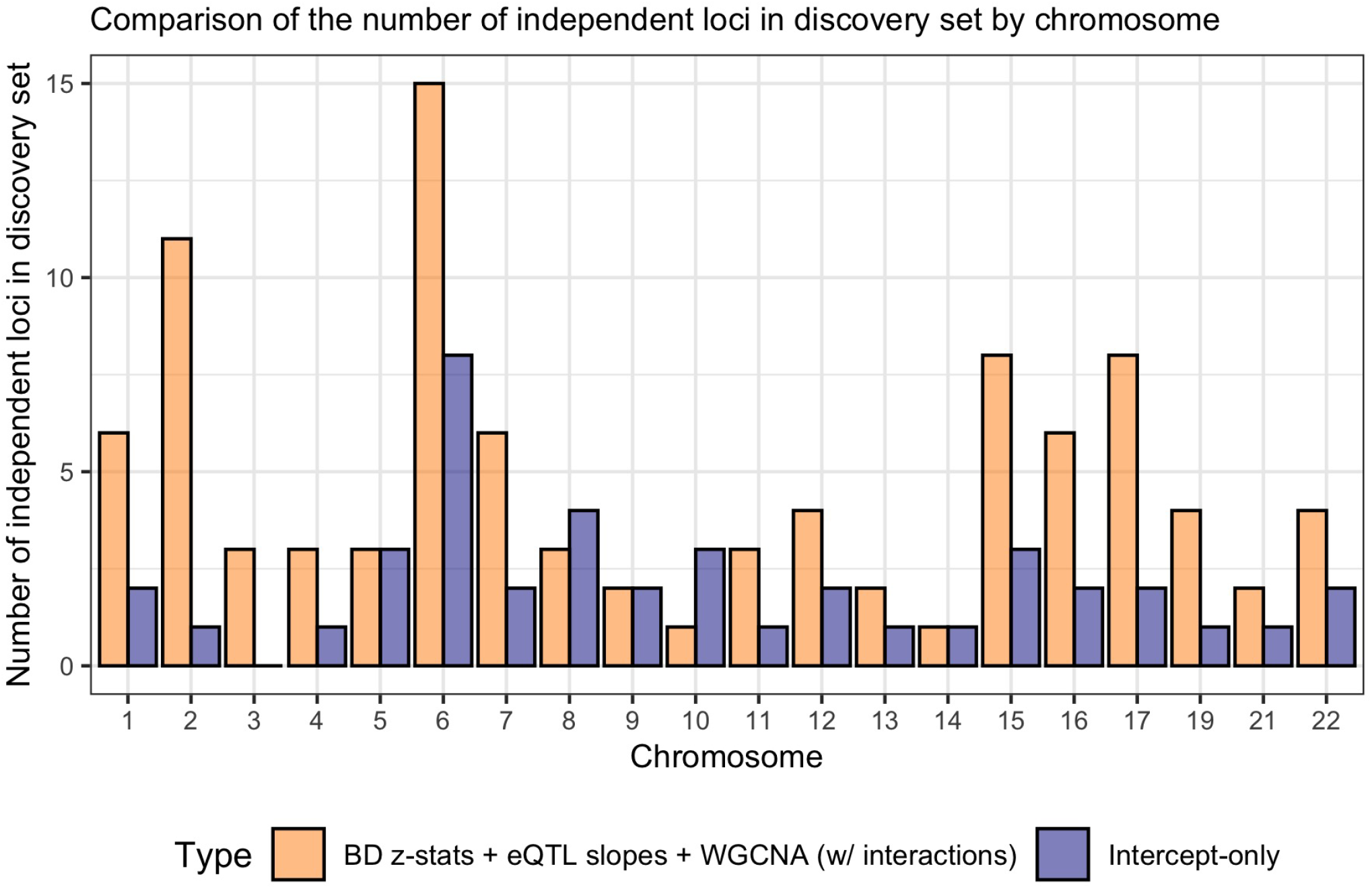
Comparison of the number of independent loci in the AdaPT discovery sets by type for each chromosome.

**Figure S4:**
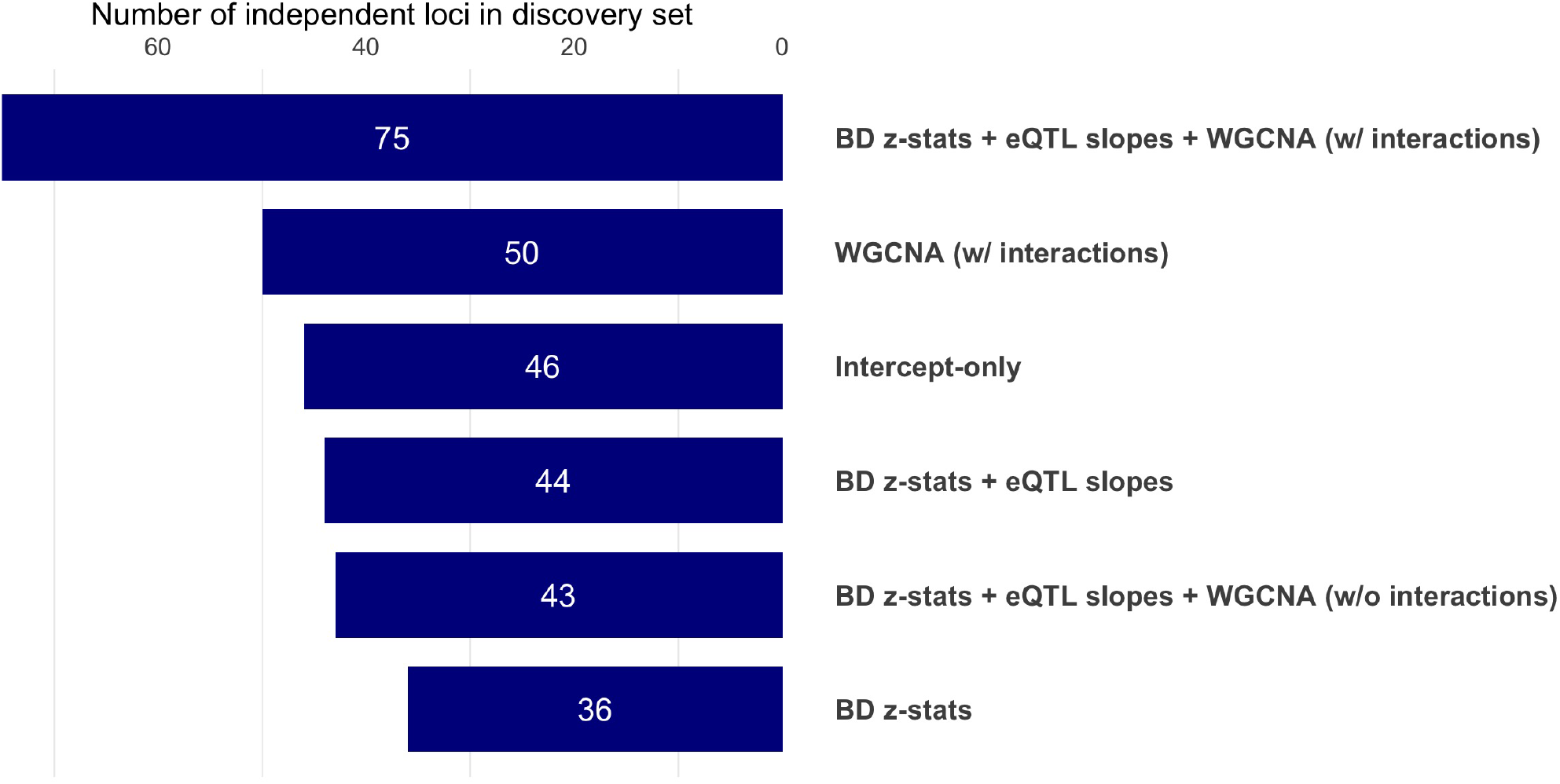
Comparison of the number of independent loci for each discovery set at target *α* = 0.05, based on LD pruning with the with *2014-only* SCZ p-values.

In Figure S8 we display the p-value distributions comparing the enrichment for membership in the different WGCNA modules reported by Werling et al. (2019). While many of the WGCNA modules lack clear evidence or contain too few eSNPs, as denoted by their respective y-axes, the *cyan* and *salmon* modules display noticeable enrichment. Additionally, as mentioned previously, membership in the *gray* module displays a lack of enrichment versus no associated cis-eQTL gene affiliated with the unassigned WGCNA module.

As additional context for the improved performance from using all covariates with interactions, Figures S9(A-B) display the change in partial dependence between the BD z-statistics and probability of being non-null *π*_1_ across the AdaPT search for the AdaPT results using (A) BD tatistics only and (B) all covariates without interactions. When compared to the results using all covariates with interactions in Figure Figure 3(B), we see that both versions of these results display relatively flat relationships near the end of the AdaPT search. This provides evidence of the importance of the interactions between other covariates and the BD z-statistics in retaining discriminatory power of the eSNPs near the end of the AdaPT search.

**Figure S5:**
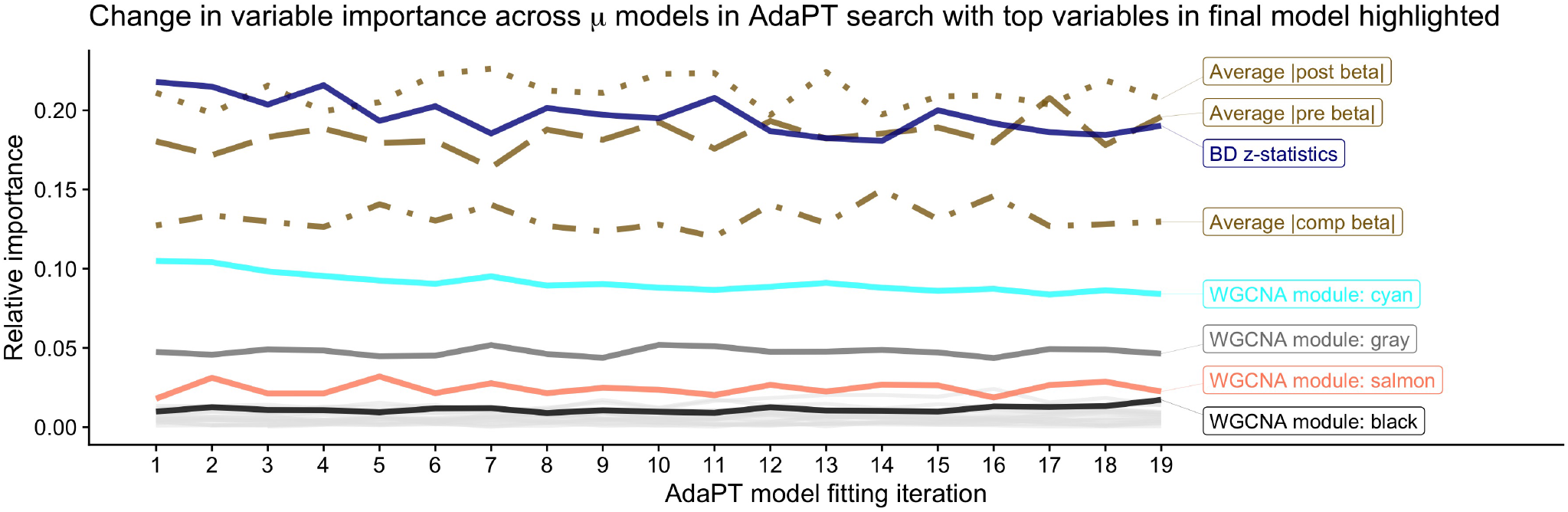
Change in variable importance for AdaPT non-null effect size *µ* model across search, with top variables in final model highlighted.

### Replication simulations

We use simulations to empirically assess the observed nominal replication rate, percentage of discoveries with p-values less than 0.05 in holdout *2018-only* studies, of 55.2% for the 843 SCZ discoveries from the *2014-only* studies at target FDR level *α* = 0.05. We use the final non-null effect size model returned by the AdaPT, 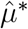, to generate simulated p-values ***p***^sim^ and nominal replication rates to compare the observed rate against. For the simulations, we assume that all 843 SCZ discoveries from the *2014-only* studies are truly non-null, and we use the actual eSNPs, their observed standard errors *σ*_14_, *σ*_18_ from the *2014-only* and *2018-only* studies respectively, as well as their actual covariates for generating ***p***^sim^. A single iteration of the simulation proceeds as follows:

- For each of the *R*_SCZ_ = 843 discoveries 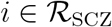:
  1. Assume test status is non-null: *H*_*i*_ = 1.
  2. Generate effect size using final AdaPT model as truth:

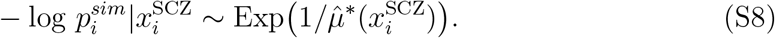
  3. Transform effect sizes to p-value 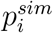.
  4. Convert simulated p-value to z-statistic 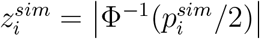.
  5. Calculate updated z-statistic to reflect observed reduction in standard error for *2018-only* studies relative to *2014-only*,

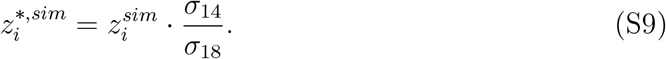
  6. Convert updated z-statistic to p-value:

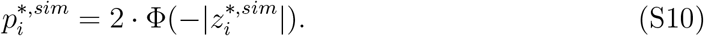
- Calculate nominal replication rate using 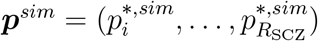

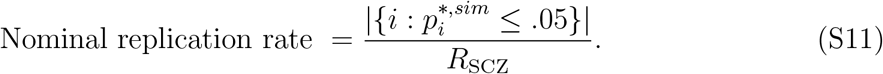

**Figure S6:**
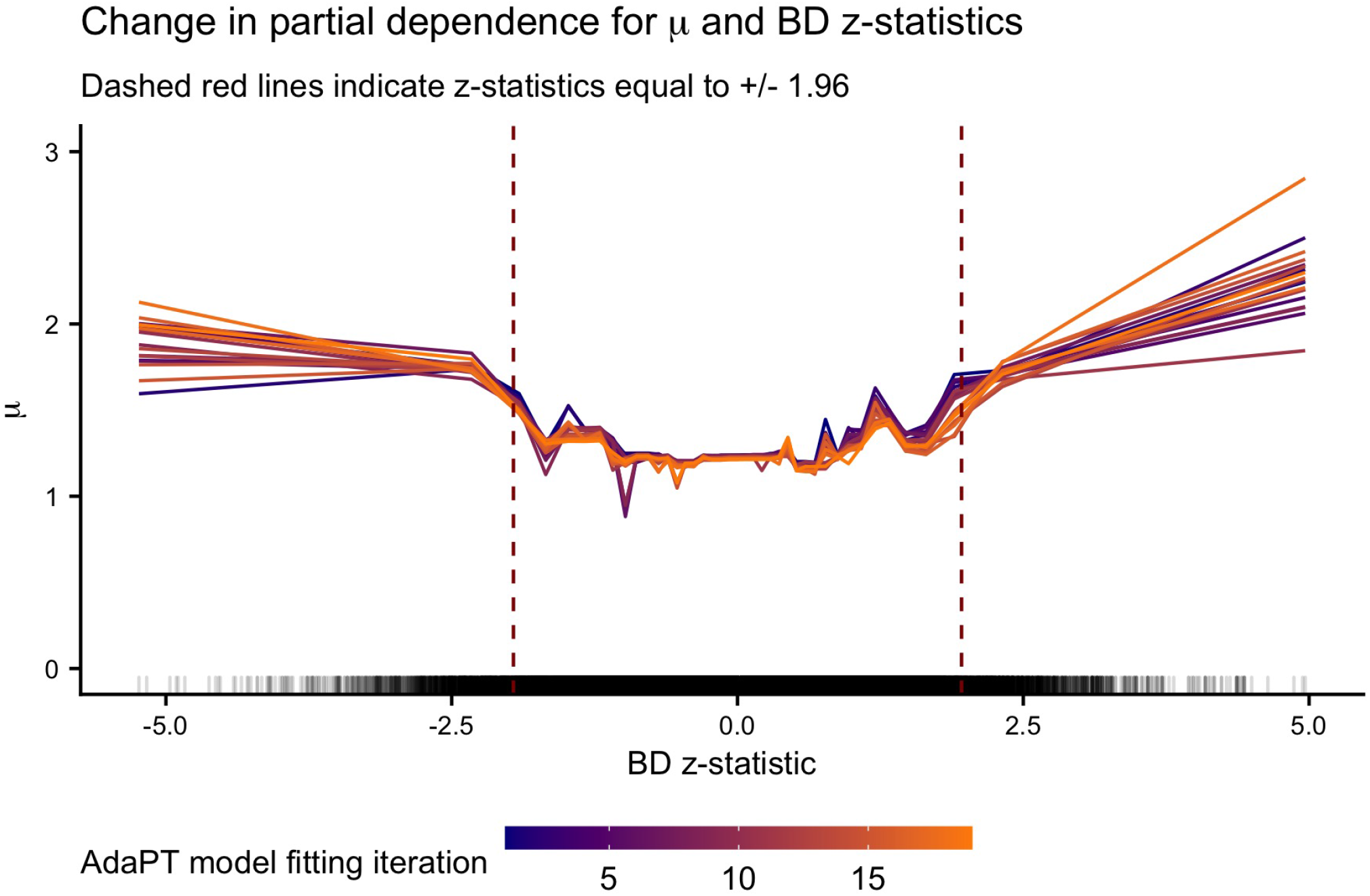
Change in partial dependence for non-null effect size *µ* and BD z-statistics across *µ* models in AdaPT search.

**Figure S7:**
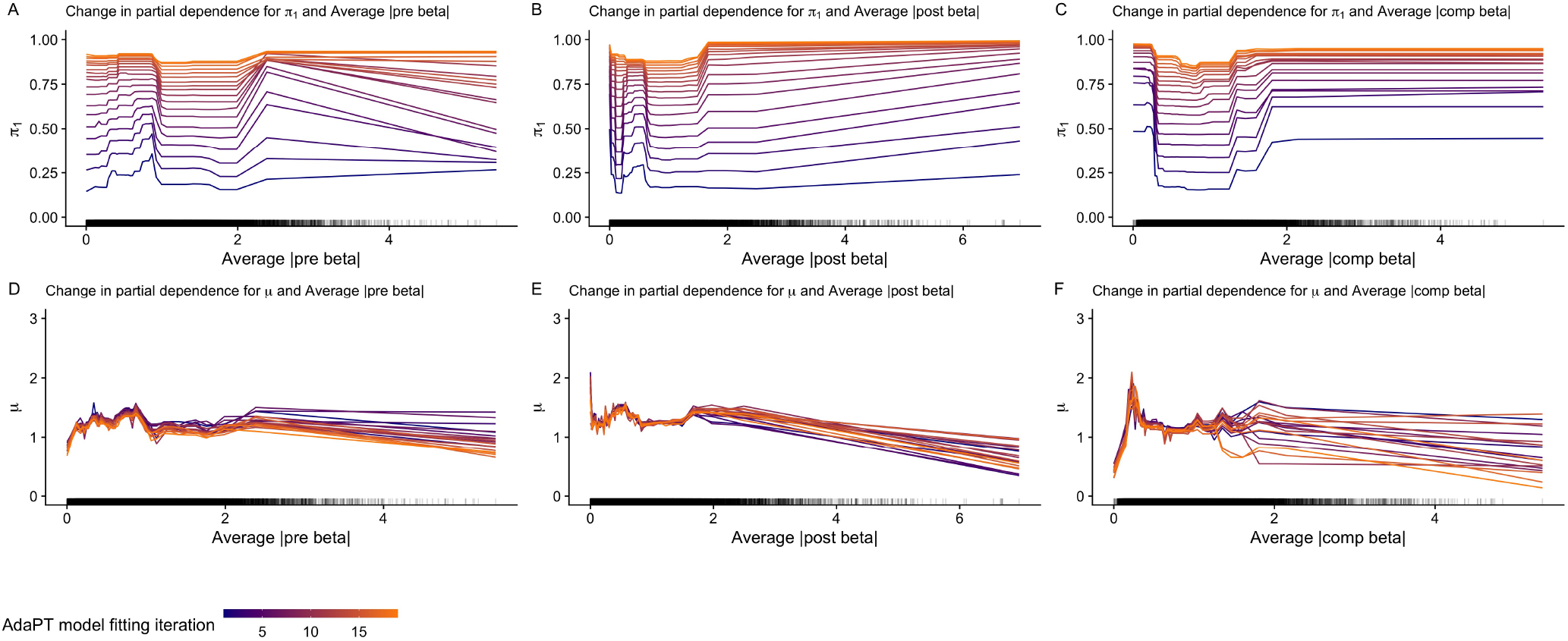
Change in partial dependence plots for probability of being non-null *π*_1_ in *(A-C)*, and the effect size under alternative *µ* in *(D-F)*, for each type of eQTL slope. Rugs along x-axis denote distribution of values for each variable.

**Figure S8:**
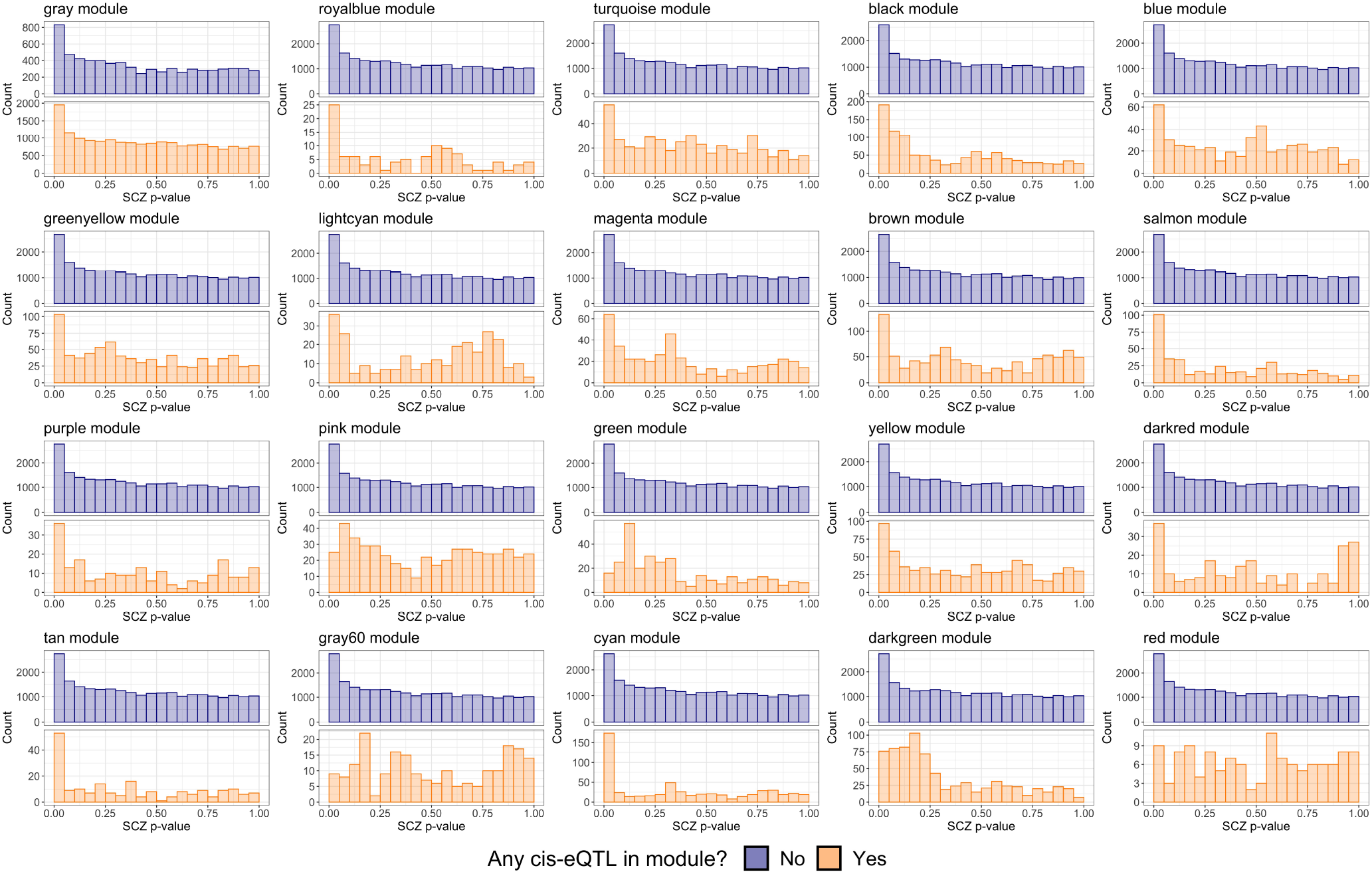
Comparison of SCZ p-value distributions from *2014* studies by whether or not the eSNP had an associated cis-eQTL gene in the module.

**Figure S9:**
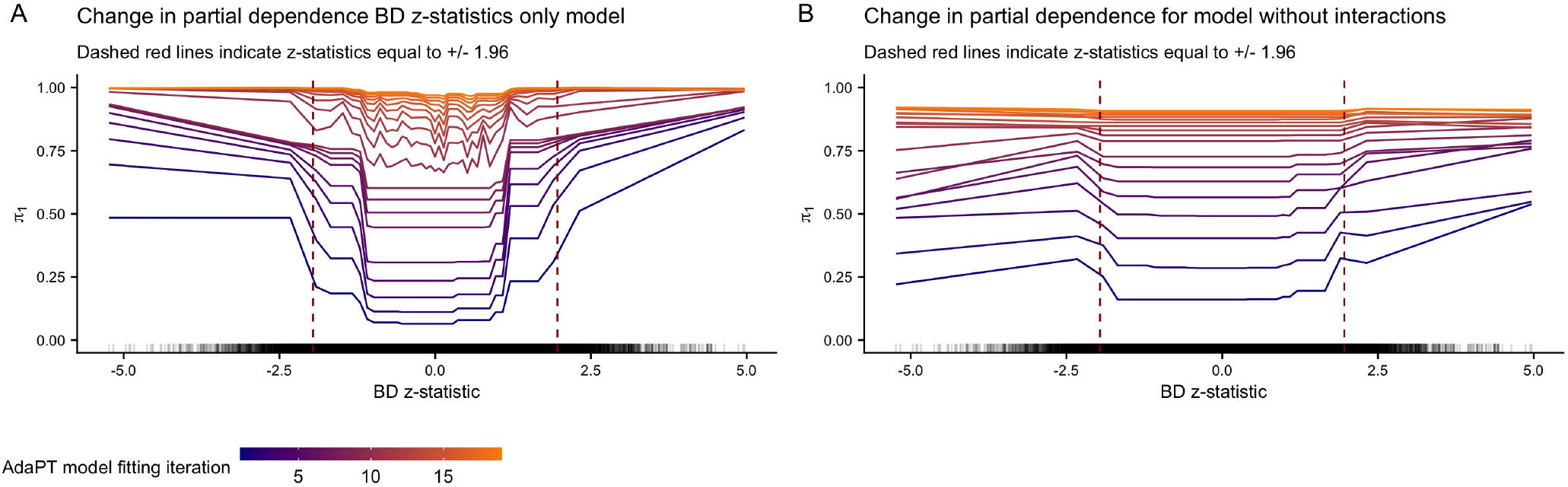
Change in partial dependence for BD z-statistics and probability of being non-null *π*_1_ for the AdaPT results using *(A)* only BD z-statistics and *(B)* all covariates without any interactions.

We repeat this process to generate ten-thousand simulated values for the nominal replication rate. The distribution of the simulated values ranges from approximately 51% to 63%, with an average and median of ≈ 57%, close to the observed rate of 55.2%. Obviously, assuming that all of the 843 rejections are truly non-null is an overtly optimistic assumption given the use of FDR error control. Thus, the average simulated nominal replication rate of ≈ 56.6% is reassuringly close to the observed rate and likely higher than what would be expected if false discoveries were accounted for among the 843 considered eSNPs.

### SCZ results with *all 2018* studies

We generate the AdaPT results using the SCZ p-values from *all-2018* studies to the same set of *n*_SCZ_ = 25, 076 eSNPs with the same covariates 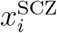. As a comparison to the results displayed in Figure 2 using the *2014-only* studies, Figures S10(A-D) display the same figures but with the results from *all 2018* at target FDR level *α* = 0.05. In contrast to before, we see that due to the increase in power from the study size, the use of modeling the auxiliary information provides a much smaller increase in power with just an approximately 19% increase in discoveries from the intercept-only results (1,865 discoveries) to using all twenty-four covariates with interactions (2,228 discoveries).

For comparison, we additionally examine the change in variable importance and partial dependence plots returned by AdaPT using *all 2018* studies. Similar to before, Figures S11(A-B) display the change in variable importance plots for both the probability of being non-null *π*_1_ and effect size under alternative *µ* models using the SCZ p-values from *all 2018* studies respectively. The results are similar to before, but with the complete sample eQTL slopes possessing the highest importance. The BD z-statistics are again highly important for *all 2018* studies, displaying the similarly increasing relationships across the AdaPT models as seen in the partial dependence plots in Figures S12(C-D). The change in partial dependence plots for the different eQTL slopes summaries are seen in Figures S13(A-F). Figure S14 displays the levels of SCZ enrichment for *all 2018* studies, revealing modules that are consistent with the *2014-only* studies such as *cyan* and *salmon*.

**Figure S10:**
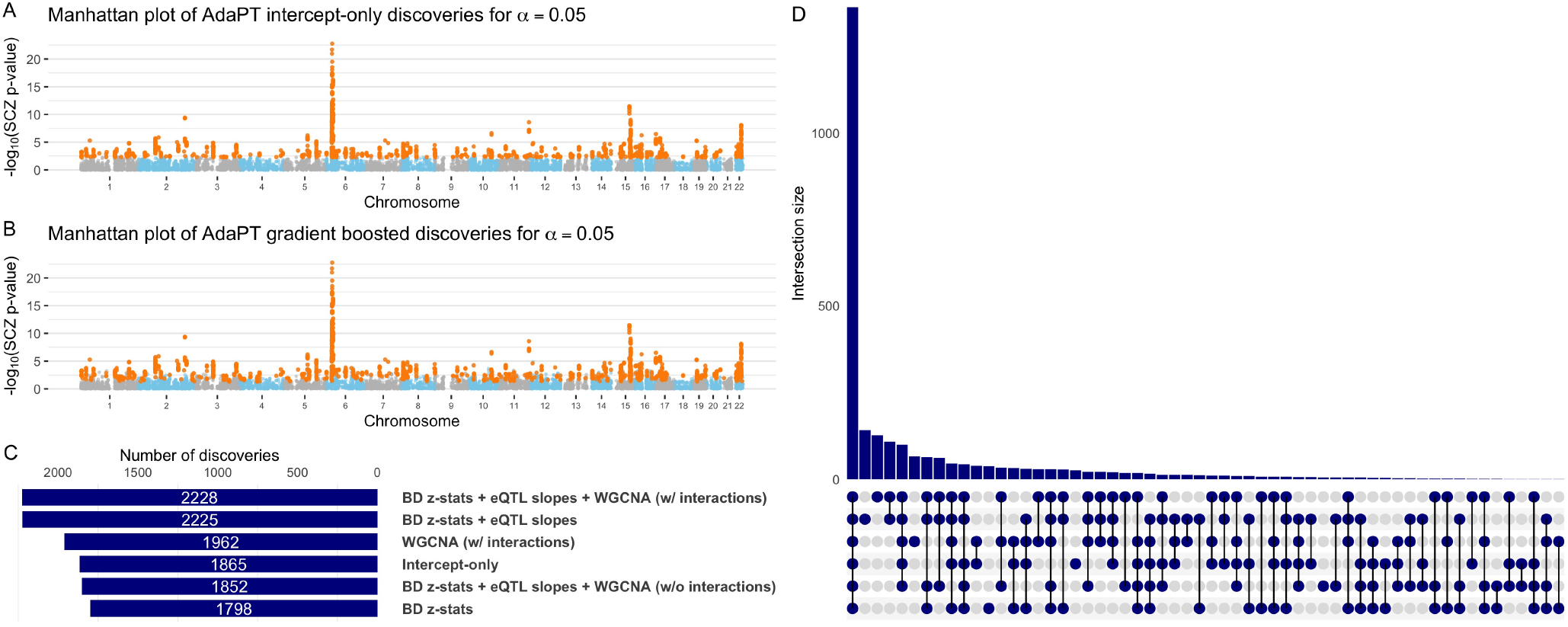
Manhattan plots of SCZ AdaPT discoveries (in orange) with *all 2018* studies using *(A)* intercept-only model compared to *(B)* covariate informed model at target *α* = 0.05. *(C)* Comparison of the number of discoveries at target *α* = 0.05 for AdaPT with varying levels of covariates and *(D)* their resulting discovery set intersections.

### Type 2 diabetes results

Using GWAS summary statistics for type 2 diabetes (T2D), unadjusted for BMI, available from Diabetes Genetics Replication And Meta-analysis (DIAGRAM) consortium (Mahajan et al. 2018), we applied our full pipeline outlined in Figure 1. Of the initial set of over twenty-three million SNPs available, we identified 176,246 eSNPs from eQTL variant-gene pairs from any GTEx tissue sample using the definition of the GTEx eSNPs explained in *Data*. Figure S15 displays the enrichment for these GTEx eSNPs compared to the original set of SNPs from the T2D GWAS results.

We create a vector of covariates 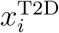 summarizing expression level information from GTEx for pancreas, liver, and two adipose tissues, *subcutaneous* and *visceral (omentum)*. Specifically, we calculate 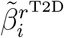 for each *r*^T2D^ in the set of tissues: pancreas, liver, adipose - subcutaneous, adipose - visceral (omentum). Additionally, we generate WGCNA module assignments using protein coding genes for pancreas samples from GTEx. To generate the WGCNA results, we only consider protein coding genes identified using the `grex` package in `R` (Xiao et al. 2018, R Core Team 2018). Additionally, all genes with expression levels of zero for over half of the provided samples were removed. This resulted in fourteen different module, including the unassigned *gray* module. Unlike the SCZ application, we do not use independent GWAS results from another phenotype.

**Figure S11:**
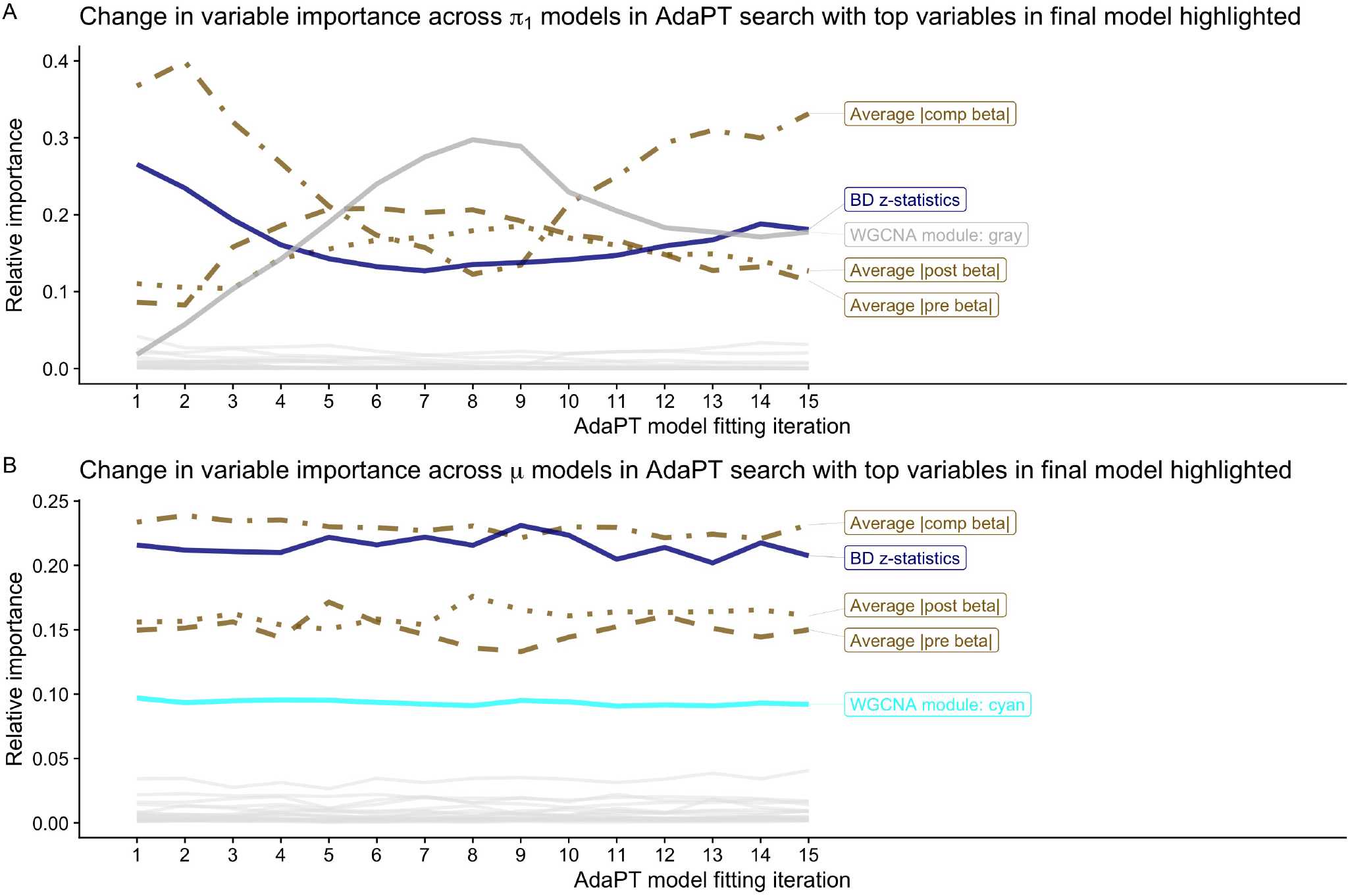
Using *all 2018* studies: change in variable importance for AdaPT *(A)* probability of being non-null *π*_1_ and *(B)* effect size under alternative *µ* models across search, with top variables in final model highlighted.

**Figure S12:**
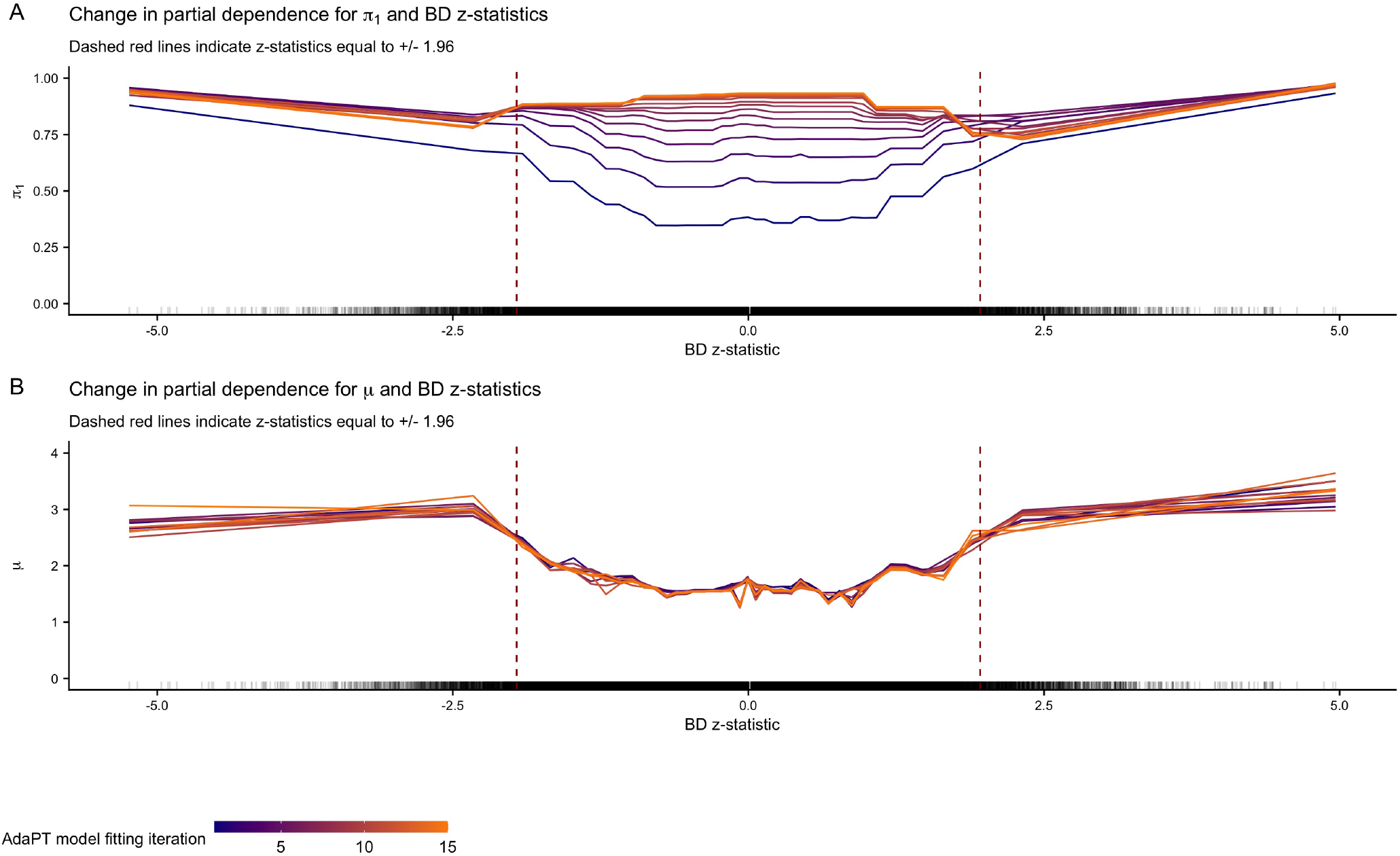
Using *all 2018* studies: change in partial dependence for BD z-statistics and AdaPT *(A)* probability of being non-null *π*_1_ and *(B)* effect size under alternative *µ* models across search.

**Figure S13:**
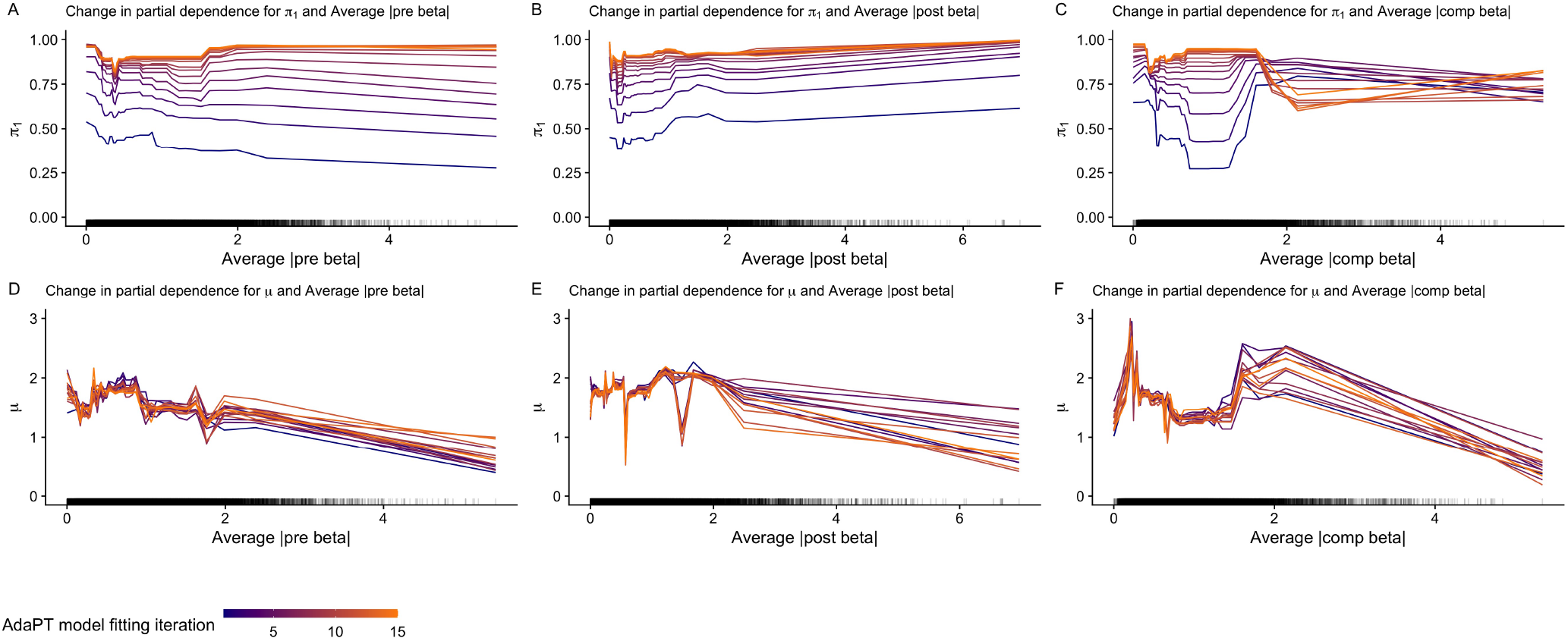
Using *all 2018* studies: change in partial dependence plots for probability of being non-null *π*_1_ in *(A-C)*, and the effect size under alternative *µ* in *(D-F)*, for each type of BrainVar eQTL slope. Rugs along x-axis denote distribution of values for each variable.

Using 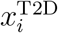 defined above, we applied AdaPT to the 176,246 GTEx eSNPs. However, we encountered an issue for this data where we were unable to discover any hypotheses at target FDR level *α* ≤ 0.05. This was due to the fact that 640 eSNPs had p-values *exactly* equal to one. While this can understandably occur with publicly available GWAS summary statistics, p-values equal to one will then *always* contribute to the *pseudo*-estimate for the number of false discoveries *A*_*t*_ during the AdaPT search (see *Methodology overview*). With a relatively high number of p-values equal to one, AdaPT is unable to search through rejection sets for lower *α* values. To overcome this challenge, we draw random replacement p-values for the 640 eSNPs from a uniform distribution between 0.97 and 1 − 1E^−15^, a value strictly less than one, to allow some leeway. We refer to this set of p-values as *adjusted*, while the original observed p-values are *unadjusted*. For comparison, Figure S16 shows the difference in the number of discoveries for the *adjusted* and *unadjusted* p-values across different target *α* values. Due to the similarity in performance for *α* values greater than 0.1, we use results for the *adjusted* p-values moving forward.

**Figure S14:**
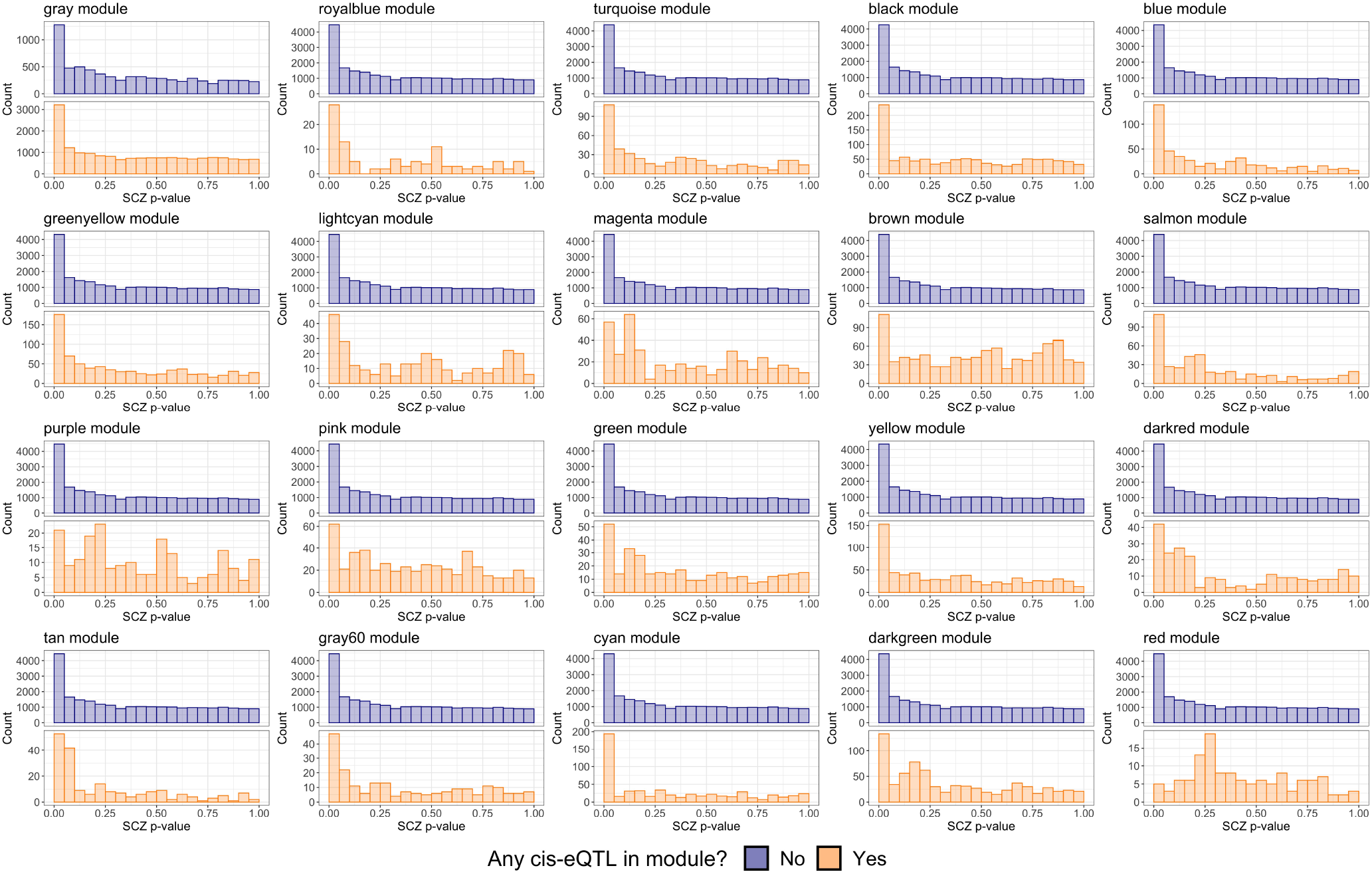
Using *all 2018* studies: comparison of SCZ p-value distributions from *2014* studies by whether or not the eSNP had an associated cis-eQTL gene in the module.

**Figure S15:**
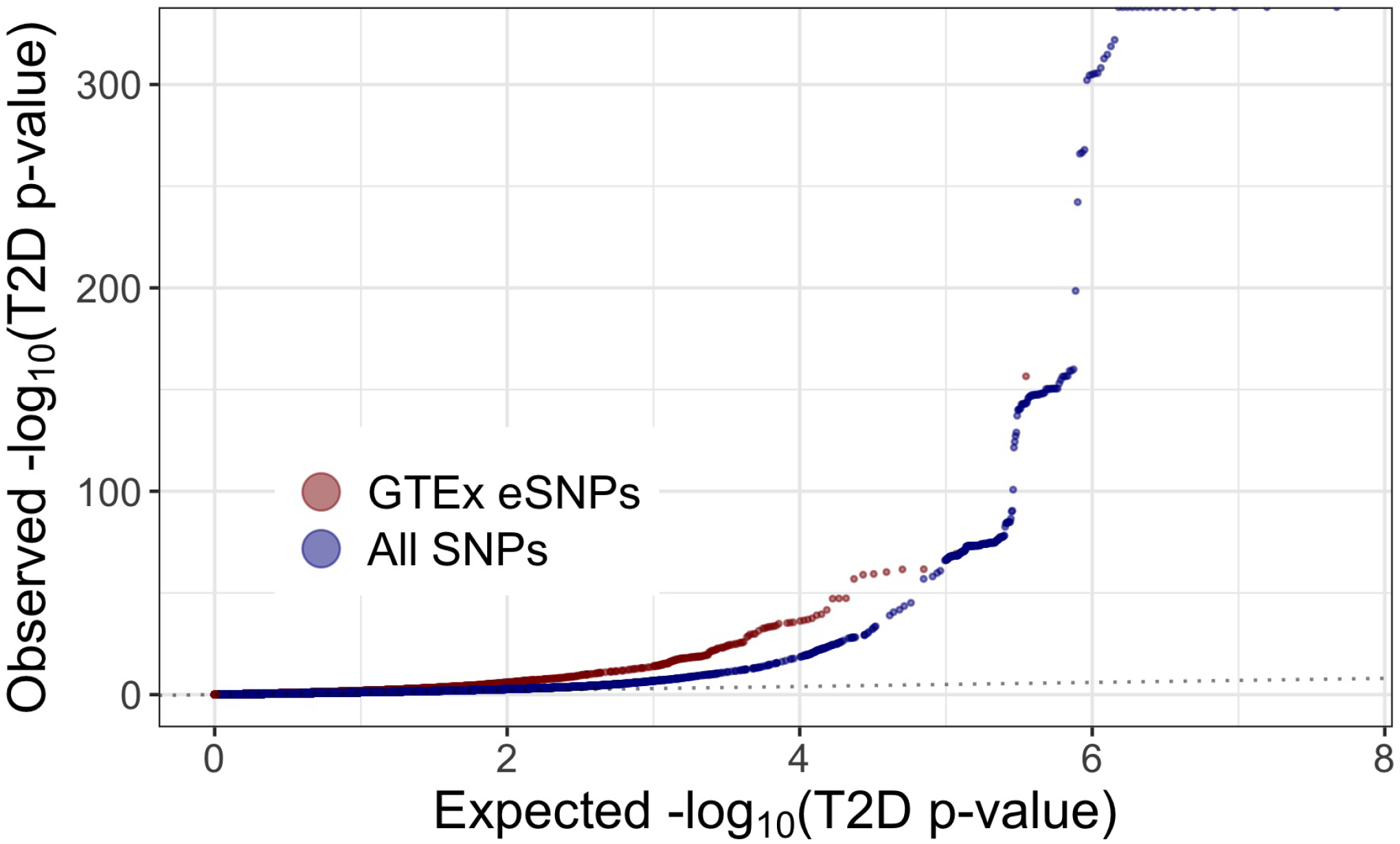
A comparison of qq-plots revealing T2D enrichment for GTEx eSNPs compared to full set of SNPs.

At target FDR level *α* = 0.05, AdaPT yields 14,920 T2D discoveries using the *adjusted* p-values with covariates 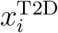 (compared to 14,693 intercept-only discoveries). The change in variable importance for the T2D AdaPT models are displayed in Figure S17. This set of eSNPs is associated with 5,970 cis-eQTL genes for which we then applied gene ontology enrichment analysis to (Ashburner et al. 2000, The Gene Ontology Consortium 2018), identifying the gene enrichment for biological processes displayed in Figure 5.

### BMI results

We also applied our pipeline of analysis to BMI, unadjusted for waist-to-hip ratio (WHR), using GWAS results for individuals of European ancestry available from the GIANT Consortium. Specifically, we approached BMI in the same manner as SCZ: apply AdaPT to GWAS results from earlier studies with a sample size of 322,154 individuals (Locke et al. 2015); then compare the nominal replication results on recently conducted studies with a sample size of approximately 700,000 individuals (Yengo et al. 2018). As before, all of the *2015-only* studies from Locke et al. (2015) were included as a subset of *all 2018* studies Yengo et al. (2018). Because both Locke et al. (2015) and Yengo et al. (2018) use the inverse variance-weighted fixed effects approach for meta-analysis, we then compute statistics for the studies exclusive to *2018-only* studies in Yengo et al. (2018). Additionally, to make this example more comparable to the SCZ use, we also use GWAS results for WHR (Shungin et al. 2015) as a covariate (analogous to BD for SCZ). Following pre-processing steps (matching SNPs across studies and effect alleles in both WHR and BMI), we identified 47,690 GTEx eSNPs from a set of nearly two million SNPs, based on the definition explained in *Data*. Figure S18 displays the enrichment for the GTEx eSNPs compared to the original set of pre-processed SNPs for the *2015-only* studies.

**Figure S16:**
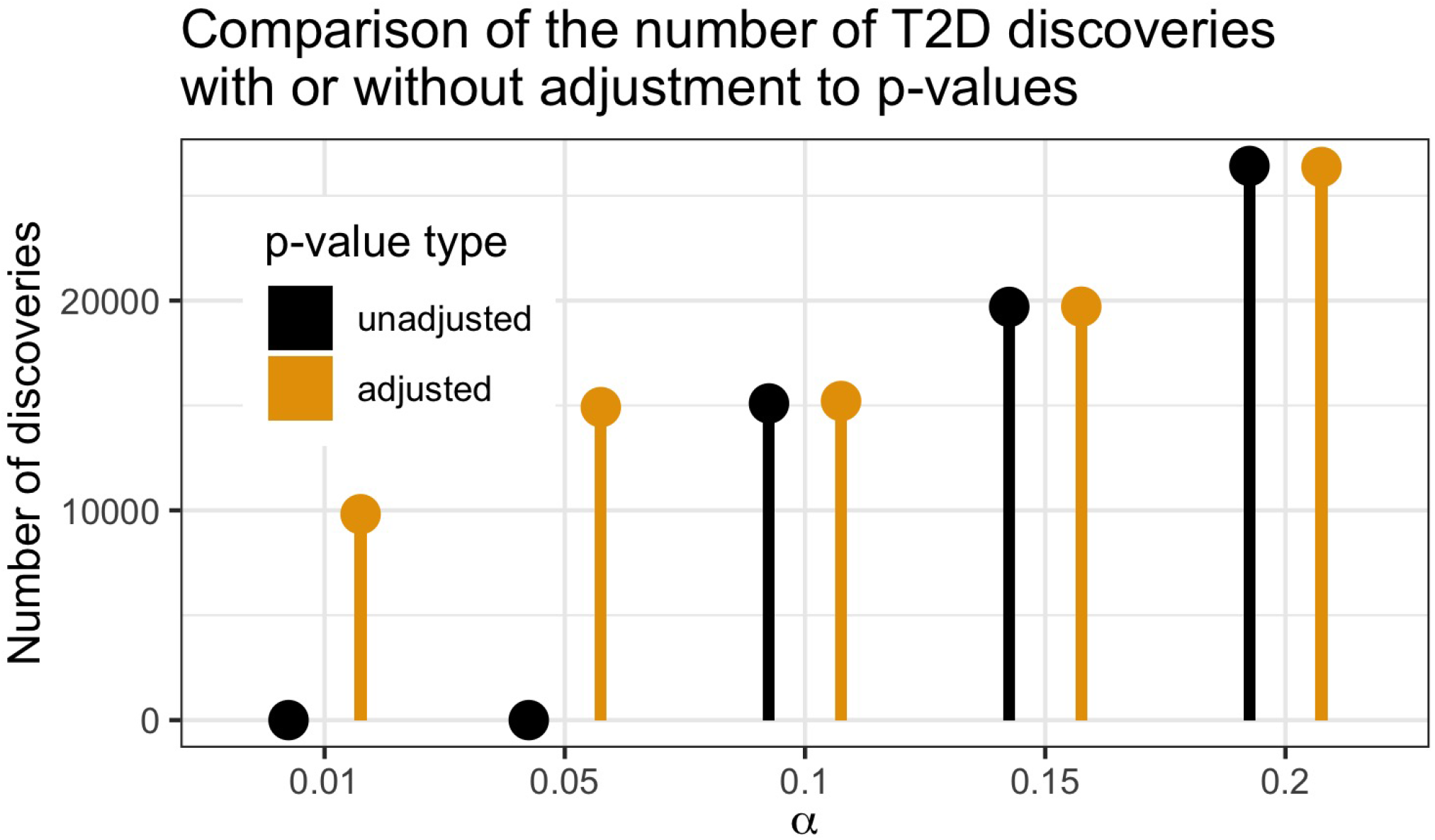
Comparison of the number of discoveries by AdaPT for T2D by whether or not the adjusted or unadjusted p-values were used.

**Figure S17:**
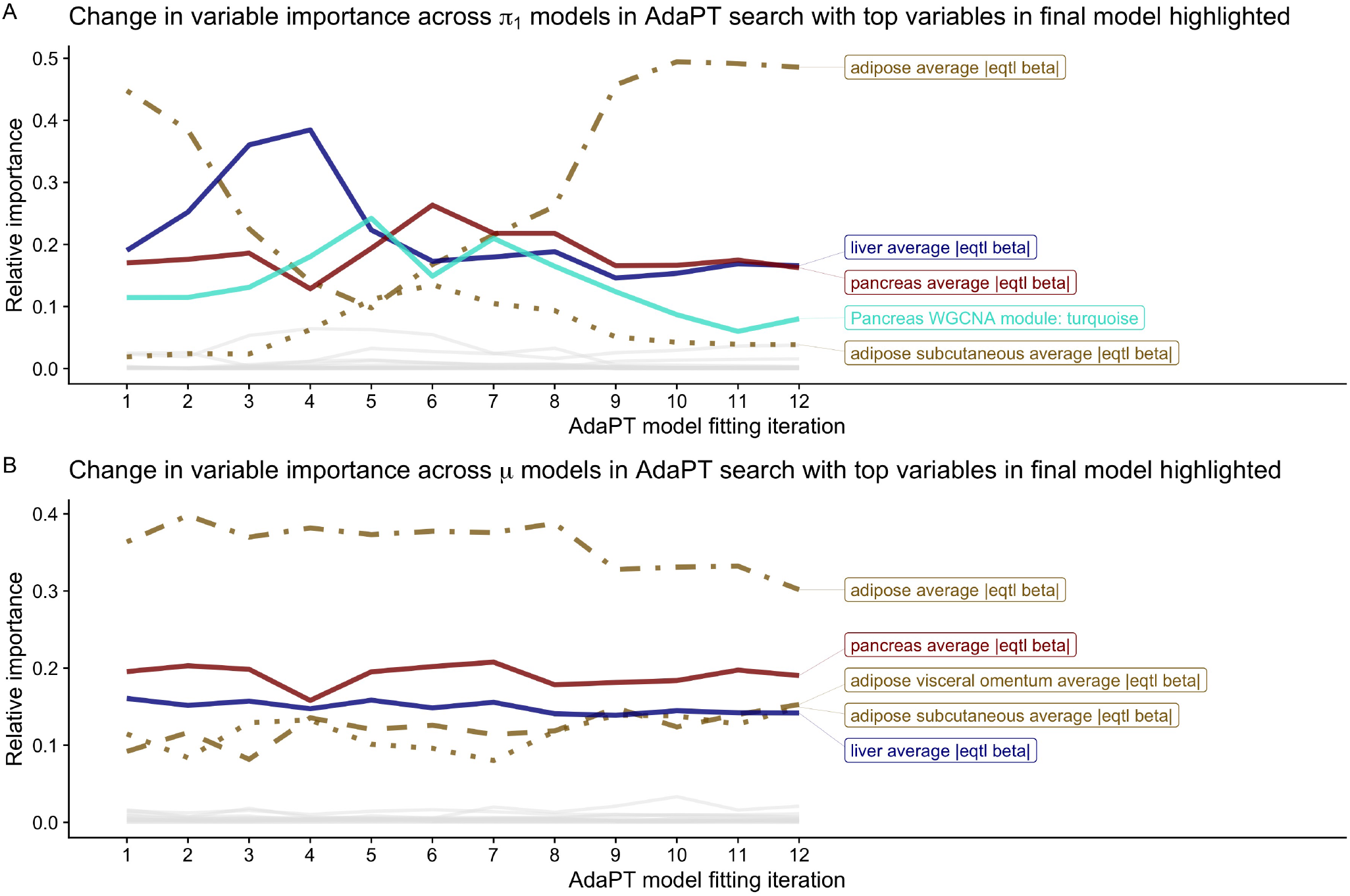
Change in T2D variable importance for AdaPT *(A)* probability of being non-null *π*_1_ and *(B)* effect size under alternative *µ* models across search, with top variables in final model highlighted.

Based on previous knowledge of BMI tissue expression associations (Locke et al. 2015), we create a vector of covariates 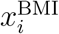 summarizing expression level information from GTEx for brain and adipose tissues (both *subcutaneous* and *visceral (omentum)*). Specifically, we calculate 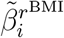 for each *r*^BMI^ ∈ {GTEx brain tissues, adipose - subcutaneous, adipose - visceral (omentum)}, where we consider the following brain tissues: (1) *amygdala*, (2) *anterior cingulate cortex BA24*, (3) *caudate basal ganglia*, (4) *cerebellar hemisphere*, (5) *frontal cortex BA9*, (6) *hippocampus*, (7) *hypothalamus*, (8) *nucleus accumbens basal ganglia*, (9) *putamen basal ganglia*, (10) *spinal cord cervical c-1*, and (11) *substantia nigra*. We do not consider the available *cerebellum cortex* tissue samples from GTEx as these are duplicates of *cerebellar hemisphere* and *frontal cortex BA9* respectively. We instead only use the samples taken the same time as the other brain sub-regions at the University of Miami Brain Endowment Bank, preserved by snap freezing (see GTEx FAQs).

We also created an aggregate across 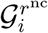, all cis-eQTL genes associated with eSNP *i* for each non-cerebellar hemisphere brain tissue region *r*^nc^,

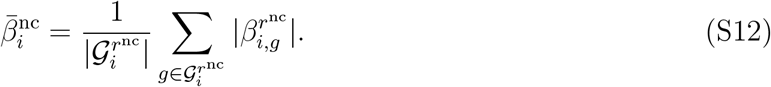

We did not include the cerebellum tissue samples in this aggregate due to the reported distinctness of the cerebellum relative to other brain tissue samples (GTEx Consortium 2015). Similarly, we computed an average across the two adipose tissues. As before, when calculating the various eQTL slopes summaries, if eSNP *i* was not an eQTL for a particular region then we impute a value of zero reflecting the lack of associated expression.

Furthermore, WGCNA module assignments were generated using protein coding genes for three different sets of tissues: (1) all non-cerebellar hemisphere brain tissues, (2) cerebellar hemisphere only tissue, and (3) adipose tissues (using same settings described previously in *Type 2 diabetes results*). Together with the WHR z-statistics and covariates accounting for the associations and WGCNA module indicators, 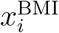 contained 110 variables.

For BMI eSPS, 376 have p-value exactly equal to one, leading to the same problem as we encountered in the T2D analysis. Again, we proceed by randomly drawing replacement p-values for these 376 eSNPs from a uniform distribution between 0.97 and 1 − 1E^−15^. Figure S19 shows how AdaPT fails to obtain any discoveries across the various *α* levels without making an adjustment to the p-values. With this limitation recognized, we proceed to focus on the discoveries returned by AdaPT using the adjusted p-values at *α* = 0.05.

Unlike SCZ and T2D, AdaPT using all of the covariates (with the same tuning parameters as SCZ) detected fewer discoveries: 1,383 eSNPs compared to 1,624 eSNPs discovered by the intercept-only AdaPT model at target FDR level *α* = 0.05. With further boosting regularization, beyond what is considered here, one could achieve the intercept-only results with gradient boosted trees. Of these 1,383 discoveries, approximately 83% (1,140 eSNPs) were nominal replications with p-values less than or equal to 0.05 in the independent *2018-only* studies. Figure S20 displays the increasing smoothing spline relationship between the *2018-only* p-values and the resulting *2015-only* q-values from the AdaPT search on the log_1_0 scale. The much higher observed nominal replication rate is not surprising given the well powered size of the BMI studies, as indicated by the y-axis of Figure S20, which reflects the level of enrichment for the *2018-only* studies.

Additionally, gene ontology enrichment analysis for the 1,383 discoveries using all covariates revealed no significant biological process enrichment at target FDR level *α* = 0.05. One concern is that a model with 110 variables is excessive, because the variable importance plots for the BMI AdaPT models in Figures S21(A-B), along with the partial dependence plots in Figures S22(A-B), emphasize the relative importance of the WHR z-statistics compared to other covariates. To test this conjecture, we explored two simpler models using (1) WHR z-statistics only and (2) WHR z-statistics with eQTL slope summaries. These produced 1,324 and 1,351 discoveries at the 0.05 level, respectively. We conclude that the available covariates do not provide sufficient additional information beyond the signal available with this immense sample and consequently including covariates in the AdaPT model does not increase the power of the procedure.

**Figure S18:**
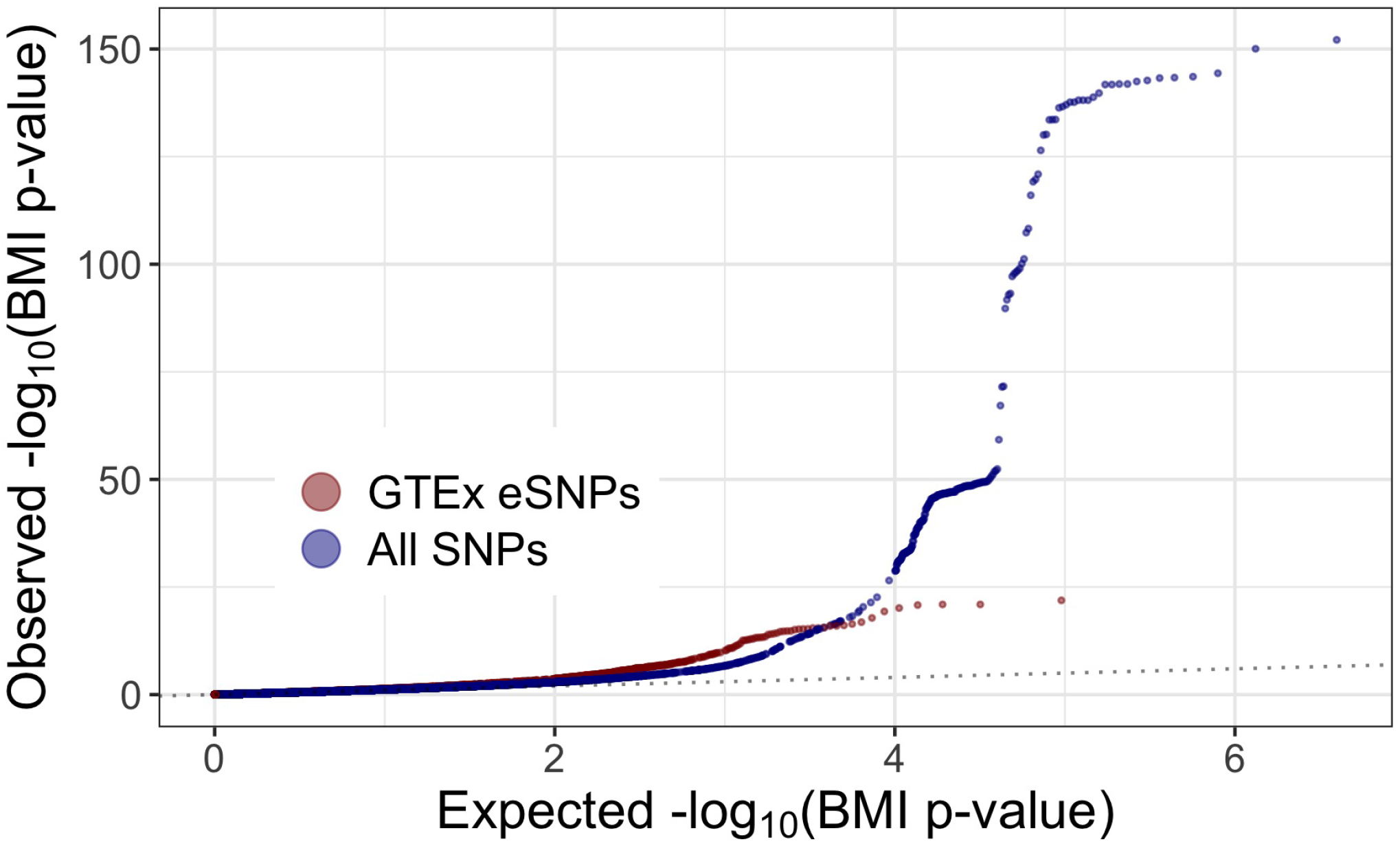
Comparison of qq-plots revealing BMI enrichment for GTEx eSNPs compared to full set of SNPs.

### CV tuning for SCZ, T2D, and BMI results

Rather than fixing the parameter settings for the XGBoost gradient boosted trees, we use the CV algorithm (detailed in *Methods*) at two steps of the search to tune the models (see the following section for justification of using two CV steps). For our search space, we evaluate a small range of values for the number of trees *P* and limit the maximum tree depth *D* to result in reasonably shallow trees (referred to as `nrounds` and `max_depth` in the `xgboost` package (Chen et al. 2019)).

First, for SCZ analysis, when exploring the improvement in discovery rate for the eSNPs by incrementally including more information, we used the following XGBoost settings:

- BD z-stats: Combinations of *P* ∈ {100, 150}, *D* ∈ {1, 6},
- BD z-stats + eQTL slopes: Combinations of *P* ∈ {100, 150}, *D* ∈ {3, 6},
- BD z-stats + eQTL slopes + WGCNA: Combinations of *P* ∈ {100, 150}, *D* ∈ {2, 3},
- WGCNA only: Combinations of *P* ∈ {100, 150}, *D* ∈ {1, 2, 3}.

**Figure S19:**
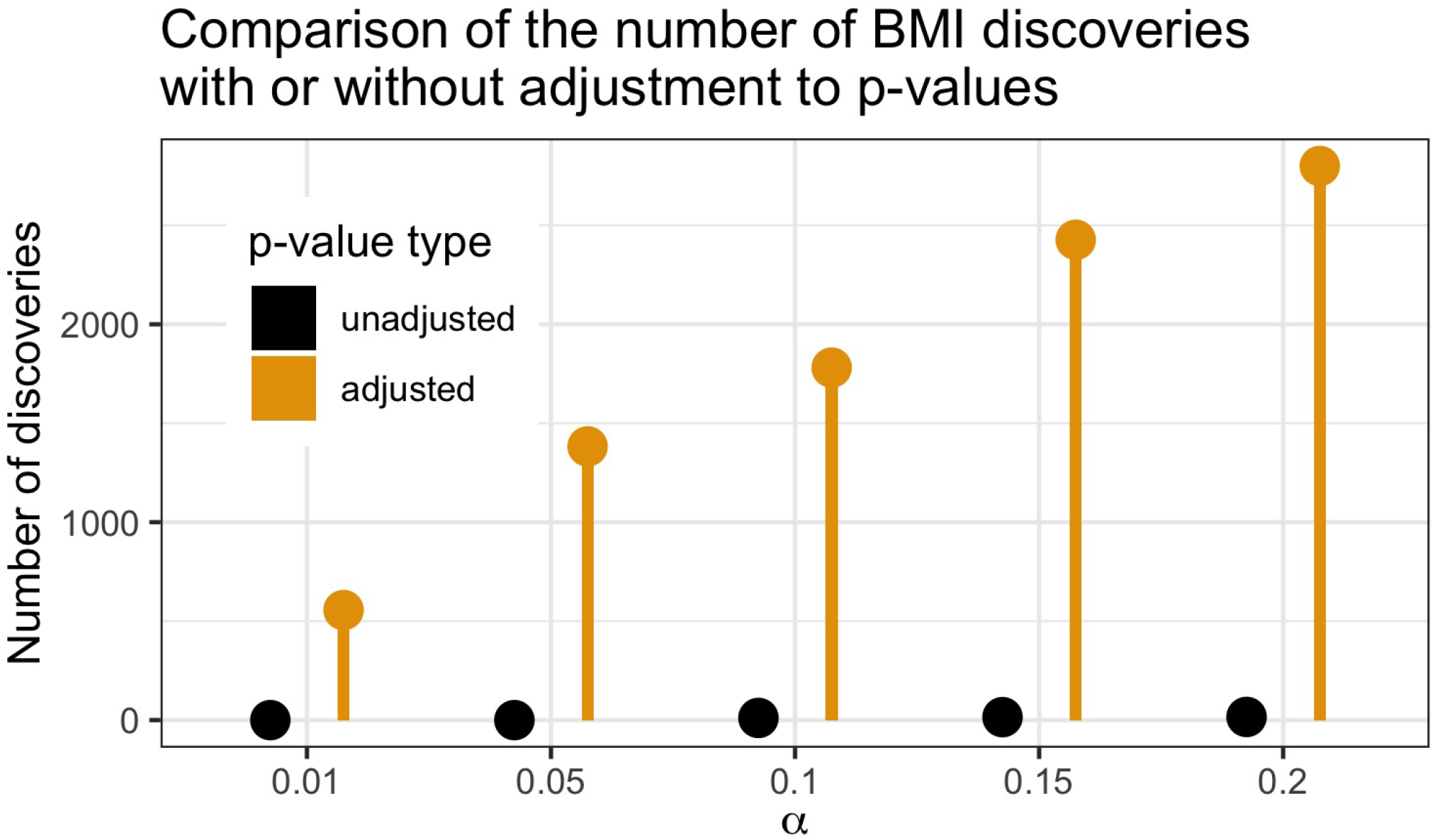
Comparison of the number of discoveries by AdaPT for BMI by whether or not the adjusted or unadjusted p-values were used.

**Figure S20:**
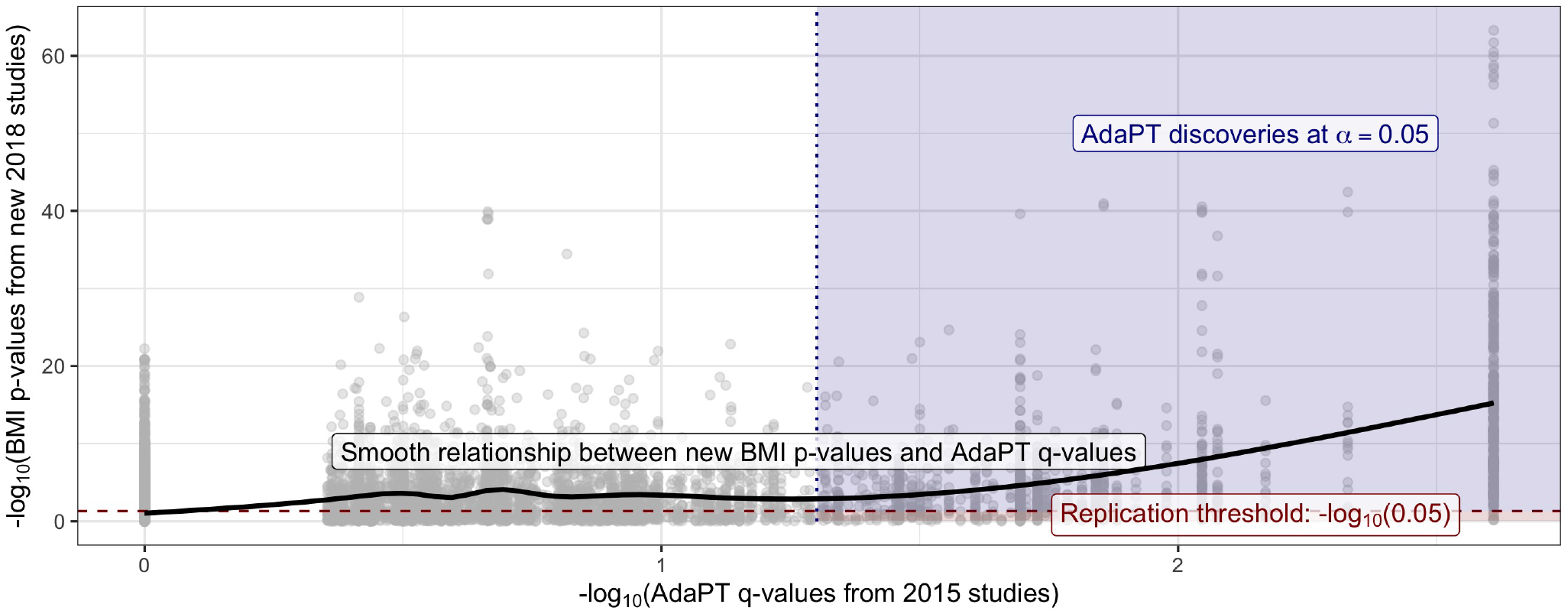
Black line displays smooth relationship between BMI p-values from *2018-only* studies and the AdaPT q-values from the *2015-only* studies. Blue-shaded region indicates AdaPT discoveries at *α* = 0.05 that are nominal replications, p-values from the *2018-only* studies < 0.05 while red denotes discoveries which failed to replicate.

We explored different settings for the different possible covariates to address the types of variables included. For instance, when using the BD z-statistics only, we considered both single-split “stumps” as well as more depth with six splits to potentially handle the variable’s symmetric relationship. Once we have all three types of covariates (BD z-statistics, eQTL slope summaries, and WGCNA results), we limit the maximum depth to be at least two to ensure possible interactions can be captured.

The selected number of trees *P* and maximum depth *D* for each of these sets of covariates is displayed in Table S1. When using only the BD z-statistics, as well as only including the eQTL slopes, the single-split settings were selected in the first CV step while the higher depth was selected in the second CV step. When using all covariates, the most complex settings (largest number of trees and largest depth) are selected in both CV steps. This agreement in selection is not surprising given the choice of the low starting threshold *s*_0_ = 0.05, which differs from the results displayed in Table S3 of the next section using *s*_0_ = 0.45. We evaluated the same possible settings for the various *all 2018* results displayed in Figures S10(C-D): the same choices for *P* and *D* displayed in Table S1 were selected in both CV steps.

**Figure S21:**
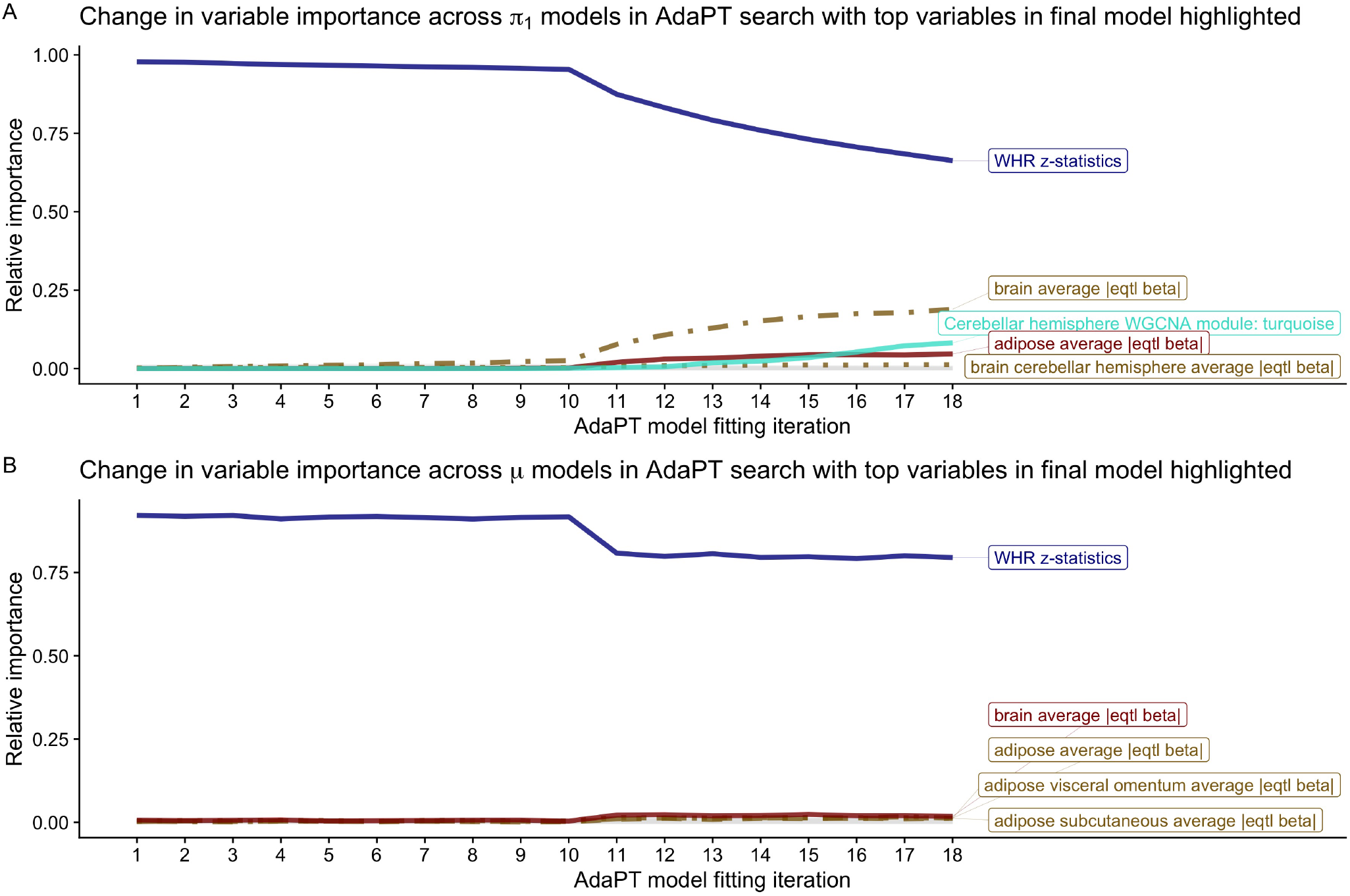
Change in BMI variable importance for AdaPT *(A)* probability of being non-null *π*_1_ and *(B)* effect size under alternative *µ* models across search, with top variables in final model highlighted.

**Figure S22:**
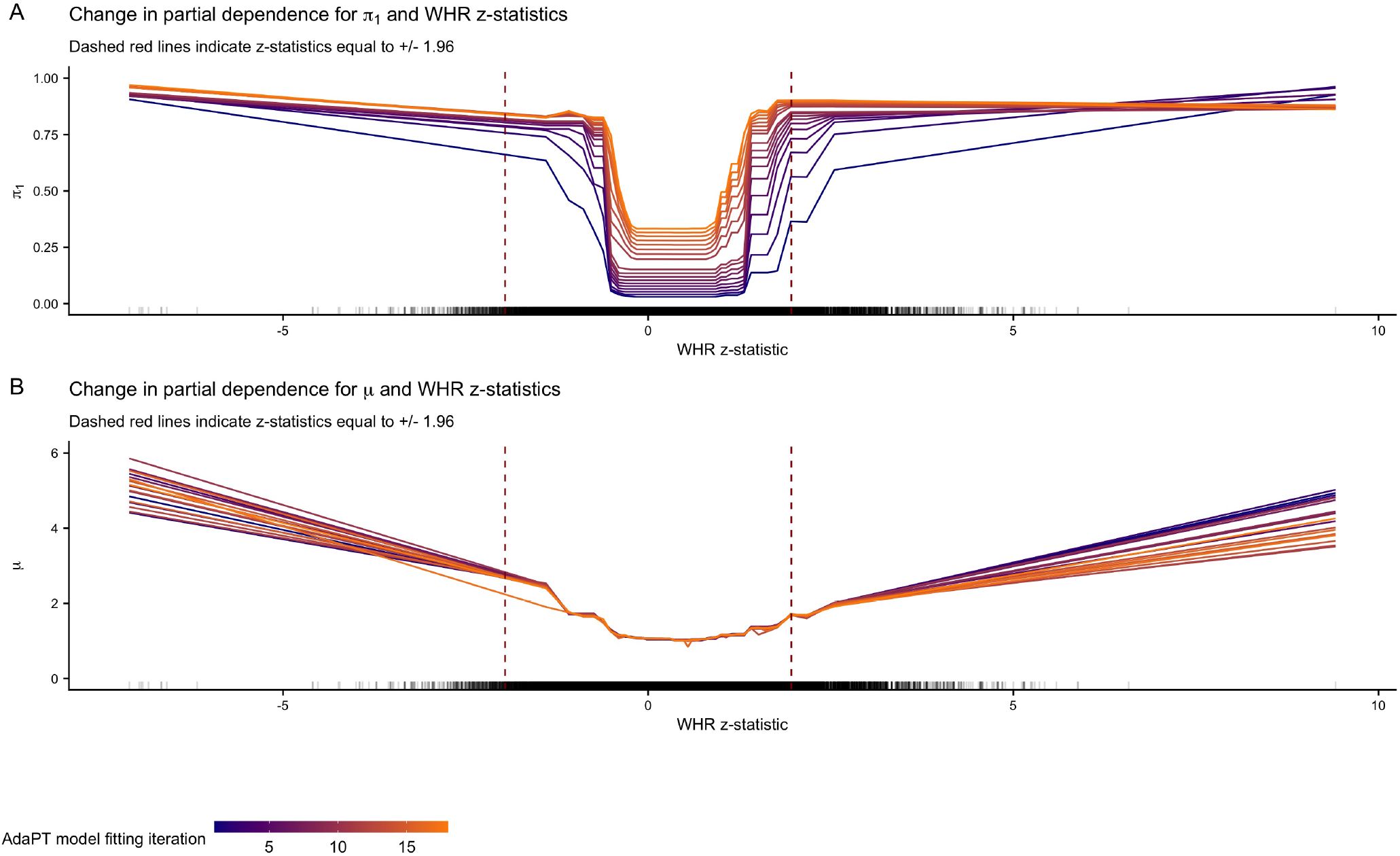
Change in BMI partial dependence for WHR z-statistics and AdaPT *(A)* probability of being non-null *π*_1_ and *(B)* effect size under alternative *µ* models across search.

For the T2D and BMI results with their full set of covariates, we evaluated four combinations: (1) *P* = 100*, D* = 2, (2) *P* = 150*, D* = 2, (3) *P* = 100*, D* = 3, and (4) *P* = 150*, D* = 3. For the BMI results using only WHR z-statistics, we varied over *P* ∈ {100, 150} and *D* ∈ {1, 6}; for the results using WHR z-statistics with the eQTL slopes, we used combinations of *P* ∈ {100, 150}, *D* ∈ {3, 6}. The selected number of trees *P* and maximum depth *D* for each of these sets of AdaPT results at both CV steps is displayed in Table S2.

**Table S1:**
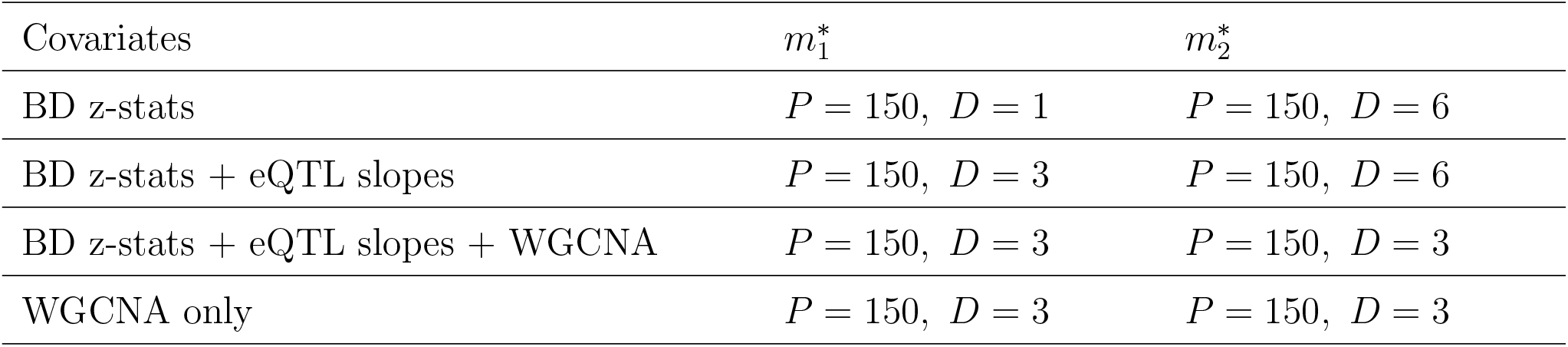
Selected boosting settings for number of trees *P* and maximum depth *D* with AdaPT CV algorithm by covariates for eSNPs in each CV step.

| Covariates | $m_1^*$ | $m_2^*$ |
| --- | --- | --- |
| BD z-stats | $P = 150$, $D = 1$ | $P = 150$, $D = 6$ |
| BD z-stats + eQTL slopes | $P = 150$, $D = 3$ | $P = 150$, $D = 6$ |
| BD z-stats + eQTL slopes + WGCNA | $P = 150$, $D = 3$ | $P = 150$, $D = 3$ |
| WGCNA only | $P = 150$, $D = 3$ | $P = 150$, $D = 3$ |

**Table S2:** Selected boosting settings for number of trees *P* and maximum depth *D* with AdaPT CV algorithm by GWAS results in each CV step.

| GWAS results | $m_1^*$ | $m_2^*$ |
| --- | --- | --- |
| T2D | $P = 100$, $D = 2$ | $P = 150$, $D = 3$ |
| BMI (all covariates) | $P = 100$, $D = 2$ | $P = 150$, $D = 3$ |
| BMI (WHR z-stats only) | $P = 150$, $D = 1$ | $P = 150$, $D = 1$ |
| BMI (WHR z-stats + eQTL slopes) | $P = 100$, $D = 3$ | $P = 150$, $D = 3$ |

**Table S3:** Selected boosting settings for number of trees *P* and maximum depth *D* with AdaPT CV algorithm by covariates for eSNPs with *s*_0_ = 0.45.

| Covariates | $m_1^*$ | $m_2^*$ |
| --- | --- | --- |
| BD z-stats | $P = 50, D = 1$ | $P = 150, D = 1$ |
| BD z-stats + eQTL slopes | $P = 100, D = 1$ | $P = 150, D = 2$ |
| BD z-stats + eQTL slopes + WGCNA | $P = 100, D = 2$ | $P = 150, D = 3$ |
| WGCNA only | $P = 150, D = 3$ | $P = 150, D = 3$ |

### Selection of *s*_0_ and number of CV steps

To justify the selection of both the starting threshold *s*_0_ and number of CV steps for the AdaPT search, we generated simulations from the first AdaPT models returned from the SCZ *2014-only* results. While these models are based on AdaPT results with a starting threshold of *s*_0_ = 0.05 following one CV step, they are only from the first model and are not explicitly parametrized by *s*_0_ and the number of CV steps. We know, however, that these first models are the result of using *P* = 150 trees with a maximum depth of *D* = 3, as indicated in Table S1 of the previous section.

Let 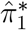 and 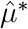 be the first models for the probability of non-null and effect size under the alternative that AdaPT returns for the eSNPs using all covariates 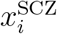. We use these models as the “truth” for generating data, in which a single iteration of the simulation proceeds as follows:

- For each eSNP 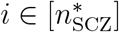
  1. Generate test status: 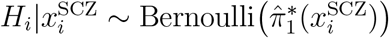.
  2. Generate imulated effect sizes:

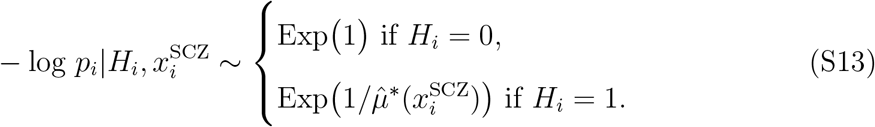
  3. Transform to p-values *p*_*i*_.
- Apply AdaPT to simulated study p-values with specified *s*_0_ and *v* CV steps with two candidate settings:
  - number of trees *P* = 100 and maximum depth *D* = 2,
  - number of trees *P* = 150 and maximum depth *D* = 3.
- Compute observed power and FDP at range of target FDR *α* values.

We generate one-hundred simulations this way for each possible threshold *s*_0_ ∈ {0.05, 0.25, 0.45} and *v* ∈ {1, 2, 5} CV steps. Figure S23 displays the average difference in power between the different starting threshold values by the number of CV steps. Although the differences are small, we see that using *s*_0_ = 0.05 results in higher power, on average, than both 0.25 and the recommended 0.45 value. Using this low starting threshold of *s*_0_ = 0.05, we then directly compute the difference in power between the different number of CV steps displayed in Figure S24. Unsurprisingly, while again the differences are small, only one CV step results in the lowest power, on average. Since the computational cost of AdaPT with CV tuning is reduced by only using two CV steps instead of a higher number, such as five, and the simulations demonstrate on average no difference in power at both *α* values of 0.05 and 0.10, we use the starting threshold of *s*_0_ = 0.05 with two CV steps in our applications of AdaPT.

In the previous section, Table S1 displayed the selections in both CV steps with *s*_0_ = 0.05. For comparison, Table S3 displays the selections using *s*_0_ = 0.45. Instead of selecting the same settings in both steps, the higher initial threshold selects the least complex settings (smallest number of trees and minimum depth) in the first CV step before flipping to the most complex settings in the second step. Intuitively, the higher initial threshold means more information is masked from the models, so it is not surprising to see less complex settings chosen. This further reinforces the use of the lower initial threshold *s*_0_ = 0.05: it starts with more revealed information and selects model settings corresponding to improved CV performance for tests with lower p-values of interest.

### Dependent p-value block simulation

To demonstrate the performance of AdaPT in the presence of dependent tests, we construct simulations with a block-correlation scheme to emulate LD structure for SNPs. We consider a setting with two independent covariates,

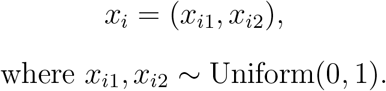

**Figure S23:**
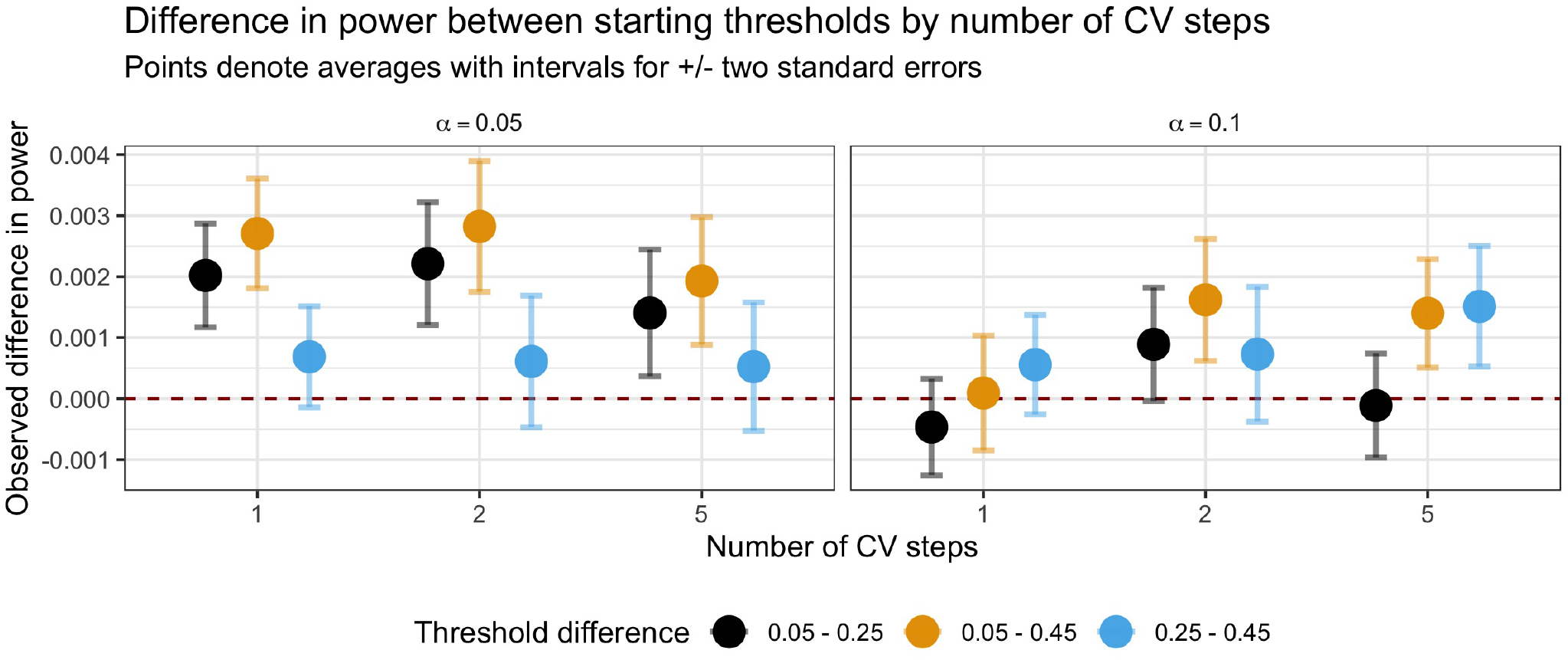
Difference in simulation power between different initial thresholds *s*_0_ for AdaPT search by number of CV steps. Points denote averages with plus/minus two standard error bars.

**Figure S24:**
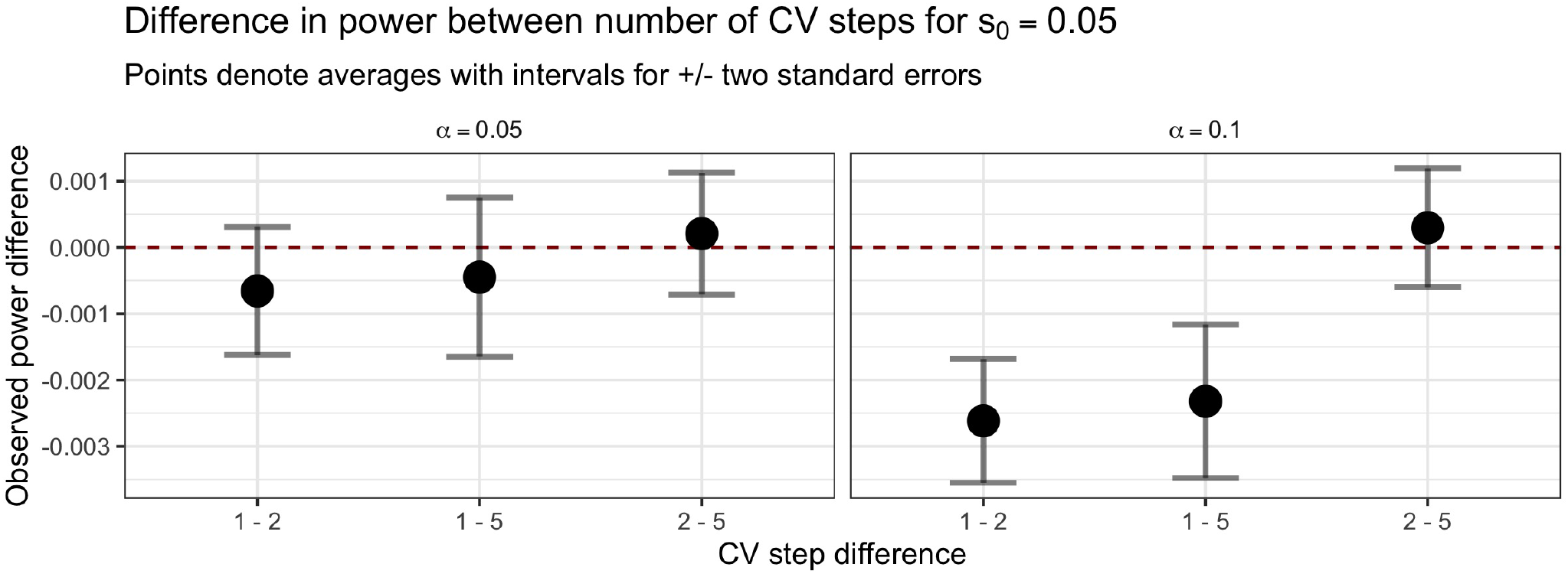
Difference in simulation power between the number of CV steps with *s*_0_ = 0.05. Points denote averages with plus/minus two standard error bars.

For each test *i ∈* [*n*], we define a linear relationship for the log-odds of being non-null using these covariates,

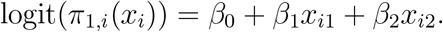

Then, the resulting status of the test *H*_*i*_ is a Bernoulli random variable based on the probability *π*_1*,i*_(*x*_*i*_) where *H*_*i*_ = 1 indicates the test *i* is non-null while *H*_*i*_ = 0 indicates a true null,

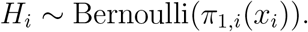

Given this test status, a vector of true effect sizes ***µ*** = *c*(*µ_i_, … , µ_n_*) is also generated as a function of the covariates,

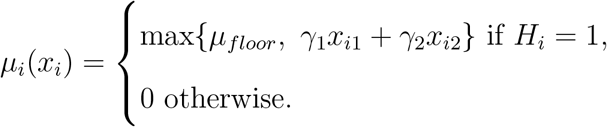

To simulate observed effect sizes, we construct an *n* × *n* covariance matrix **Σ** with *B* blocks of equal size 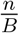. Each block *b* ∈ [*B*] has constant correlation *ρ* between all tests *within* the block, while each block is independent of each other. This results in constructing individual block covariance matrices, Σ_*b*_, with ones along the diagonal and *ρ* for the off-diagonal elements. Each of these individual matrices are placed along the diagonal of **Σ**, with the remaining off-diagonal elements set to zero so blocks are independent of each other. As an example, if each block contained only two tests they would be constructed in the following manner,

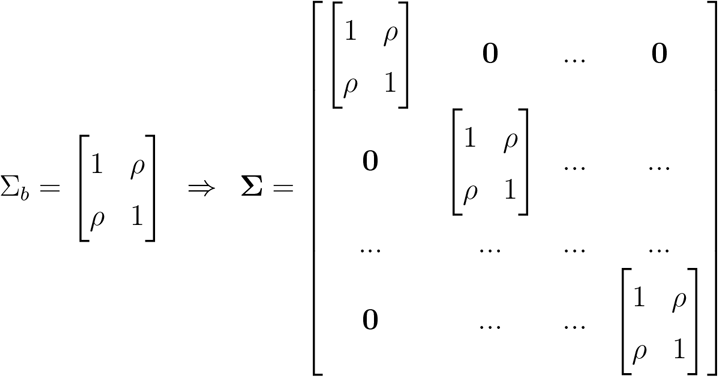

Using this block-wise construction of the covariance matrix, we then proceed to generate the vector of observed effect sizes ***z*** = (*z*_*i*_, … , *z*_*n*_) from a multivariate Gaussian distribution,

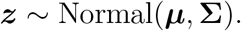

We compute the resulting two-side p-value *p*_*i*_ = 2 · Φ(−|*z*_*i*_|) for each test’s observed effect size.

For each dataset generated using this process above, we compute both the observed FDP and power for the classical BH procedure and two different versions of AdaPT:

- intercept-only,
- gradient boosted trees with covariates: *x*_*i*_ = (*x*_*i*1_, *x*_*i*2_).

We fix both *n* = 10,000 and *B* = 500 blocks, resulting in 500 blocks of twenty tests each. Rather than force all non-nulls together in the same blocks, we first calculate the minimum number of blocks required to hold all non-null tests, 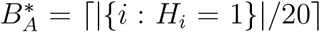. The non-null tests are then randomly assigned to 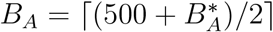 blocks, ensuring that there will be blocks containing both null and non-null tests. The |{*i* : *H*_*i*_ = 0}| tests are randomly assigned to available spots within the *B*_*A*_ blocks as well as the remaining 500 − *B*_*A*_ strictly null blocks.

In our simulations, we fix *β*_0_ = −3 and require that both *β*_1_ = *β*_2_ and *γ*_1_ = *γ*_2_. We vary the following settings in our simulations:

- block correlation *ρ* ∈ {0, 0.25, 0.5, 0.75, 1} where each block has the same value for *ρ*,
- *β*_1_, *β*_2_ ∈ {1, 2, 3},
- *µ*_*floor*_ ∈ {0.5, 1, 1.5},
- *γ*_1_, *γ*_2_ ∈ {0.5,.75, 1}.

We generate 100 simulations using the data generating process above, computing both the FDP and power for BH and the two different versions of AdaPT. For the covariate-informed version of AdaPT, we use gradient boosted trees via XGBoost with *P* = 100 trees and maximum depth *D* = 1. For both versions of AdaPT results, we start with the initial threshold of *s*_0_ = 0.45 and update the model ten times throughout the search (rather than the recommended twenty for computational speed).

Figures S25, S26, and S27 display points for the average observed FDP and power across the 100 simulations with plus/minus two standard errors bars for *µ_floor_* =0.5, 1, and 1.5 respectively, with target FDR level *α* = 0.05. The columns in each figure correspond to the different values considered for *γ*_1_ = *γ*_2_, while the rows correspond to *β*_1_ = *β*_2_. The x-axis for the figures displays the increasing block correlation *ρ*. Regardless of the simulation setting, we see that the AdaPT results when accounting for covariates (*x*_*i*1_, *x*_*i*2_) maintains valid FDR control at 0.05 similar to BH. This holds in the settings with greater effect sizes, as well as when the covariate information displays the best performance in terms of observed power (the bottom right panels of each figure). We can see that the intercept-only approach fails to achieve FDR control under block settings with perfect correlation, while the use of covariate information appears to inhibits such behavior. Our focus on positive correlation values is synonymous with the setting faced in genomics regarding LD structure. Further exploration of AdaPT’s performance in settings with arbitrary dependence structure presents an opportunity for future work, as well as accounting for covariate information that predict observed correlated noise.

### Simulations demonstrating effects of overfitting

It is possible that flexible methods like gradient boosted trees can be overfit, especially on small data sets. This could potentially lead to concerns about their incorporation in AdaPT. To assess the effects of overfitting the gradient boosted trees in AdaPT, we constructed simulated datasets using the first models returned by AdaPT on the SCZ GWAS results, 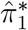 and 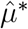, with the actual covariates 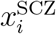 for each of the 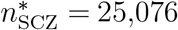 eSNPs. We then simulated data using these models in *i* SCZ the same manner previously explained for choosing *s*_0_ and the number of CV steps, and computed the observed power and FDP over a range of number of trees *P* ∈ {100, 300, 500, 700, 900}.

Figure S28(A) displays the distributions for fifty simulations of the observed FDP as the number of trees in the gradient boosted model increases. Regardless of the number of trees, we still maintain valid FDR control. However, Figure S28(B) shows as the number of trees increases, the method will overfit, resulting in a reduction in power. This reinforces that, although good model tuning can be important for power, the AdaPT method continues to maintain FDR control even as the model breaks down.

**Figure S25:**
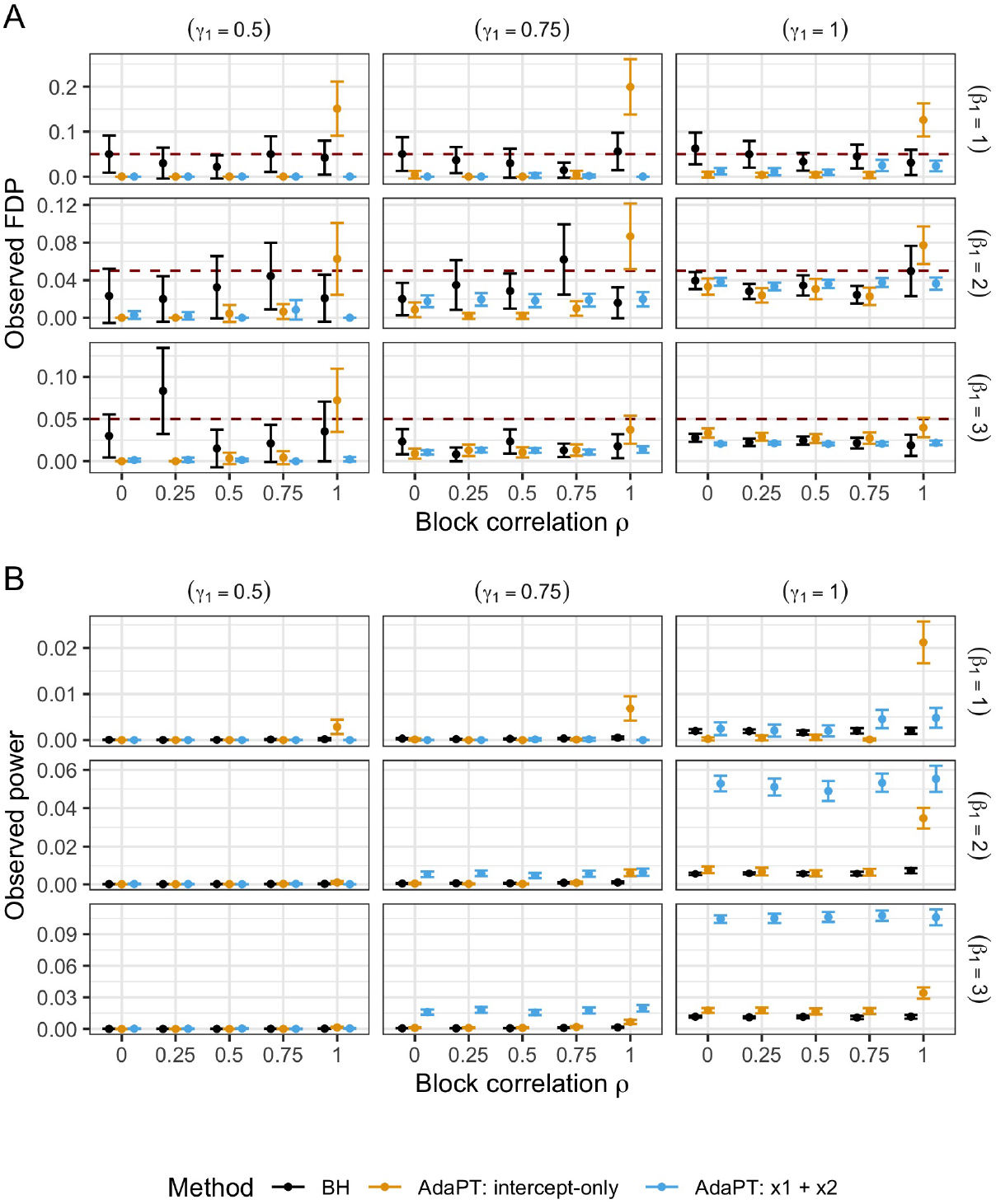
Comparison of average (A) FDP and (B) power with plus/minus two standard error bars for 100 simulations with *µ*_*floor*_ = 0.5, and varying values for *β*_1_ (rows) and *γ*_1_ (columns) and block correlation *ρ*.

**Figure S26:**
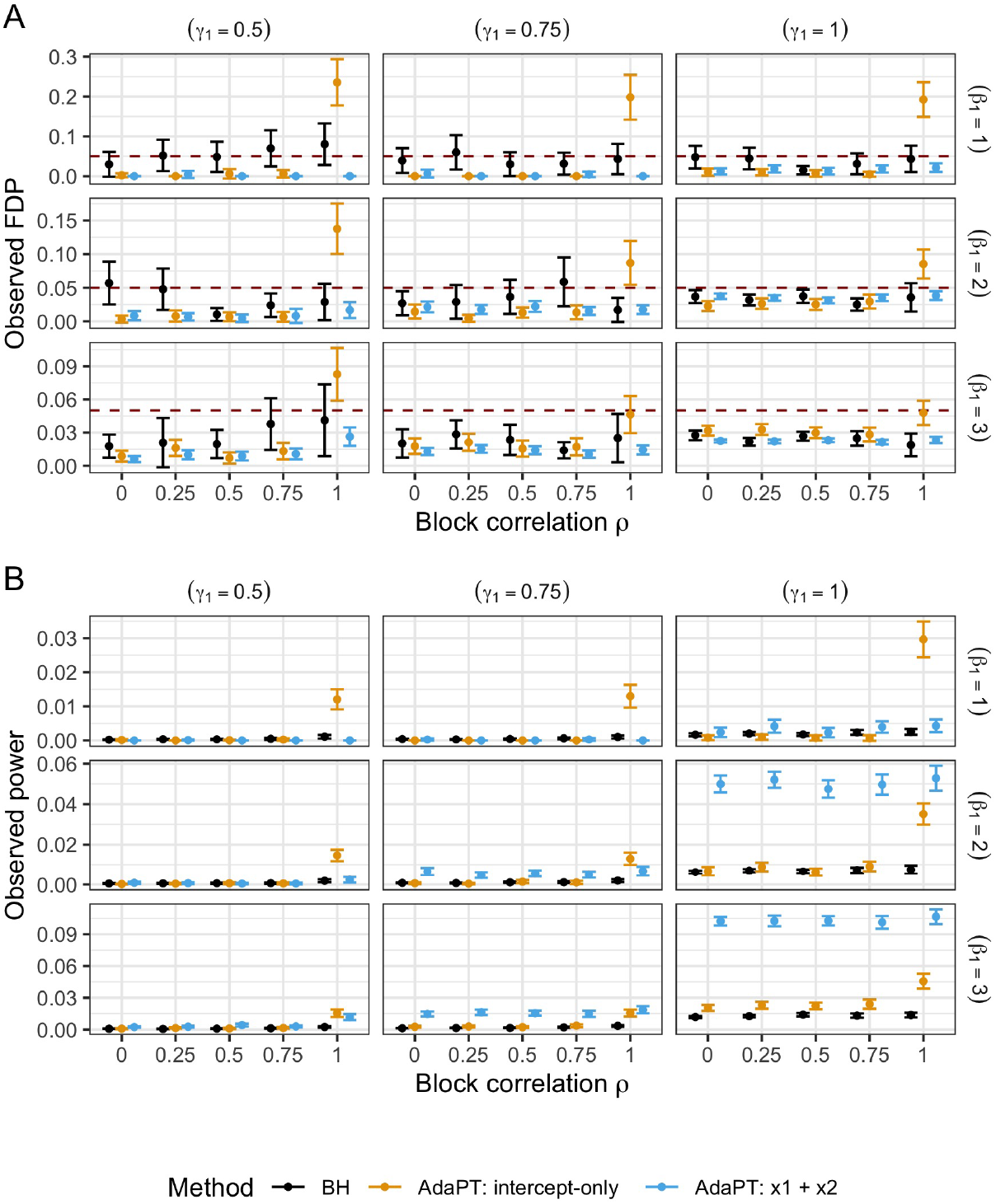
Comparison of average (A) FDP and (B) power with plus/minus two standard error bars for 100 simulations with *µ*_*floor*_ = 1, and varying values for *β*_1_ (rows) and *γ*_1_ (columns) and block correlation *ρ*.

**Figure S27:**
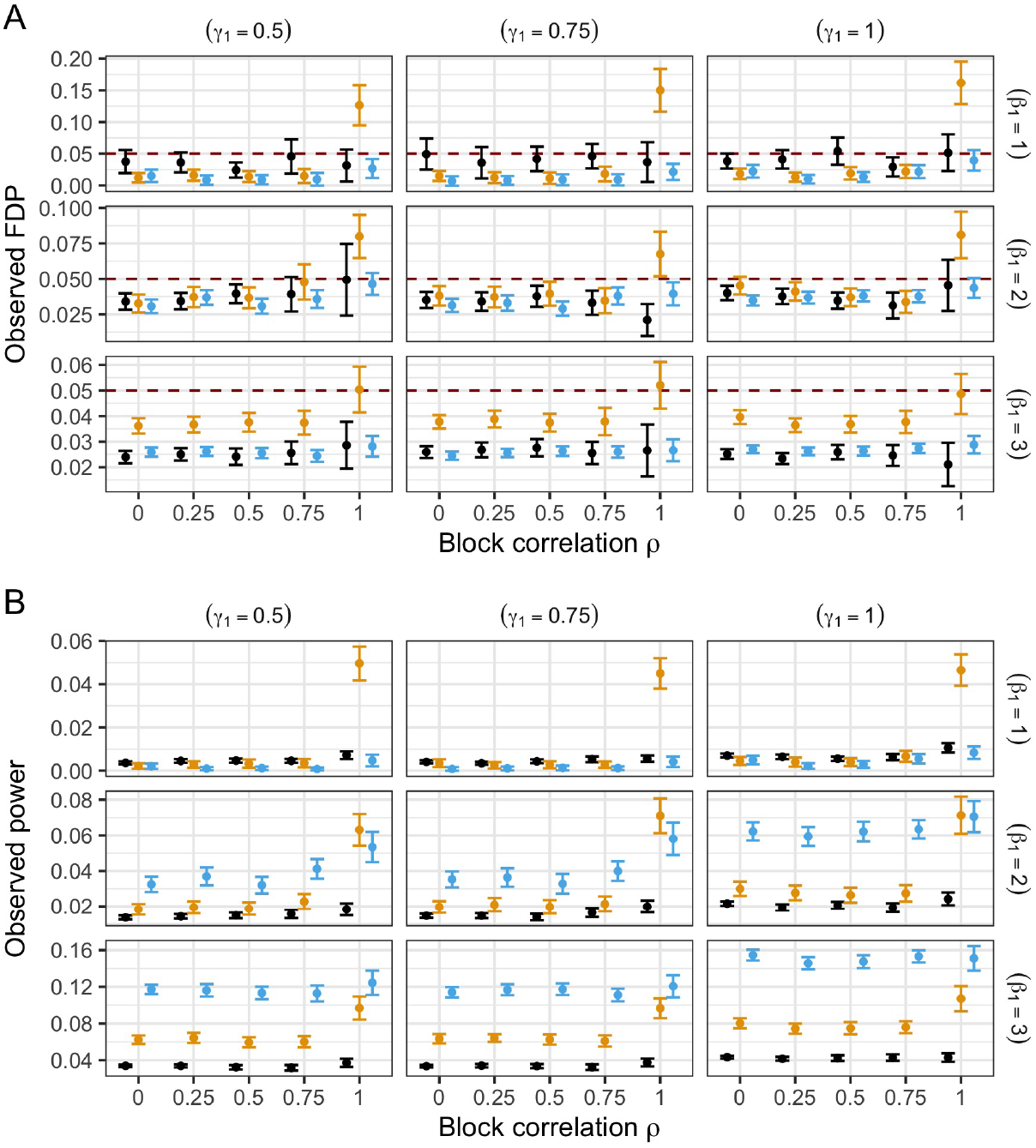
Comparison of average (A) FDP and (B) power with plus/minus two standard error bars for 100 simulations with *µ*_*floor*_ = 1.5, and varying values for *β*_1_ (rows) and *γ*_1_ (columns) and block correlation *ρ*.

**Figure S28:**
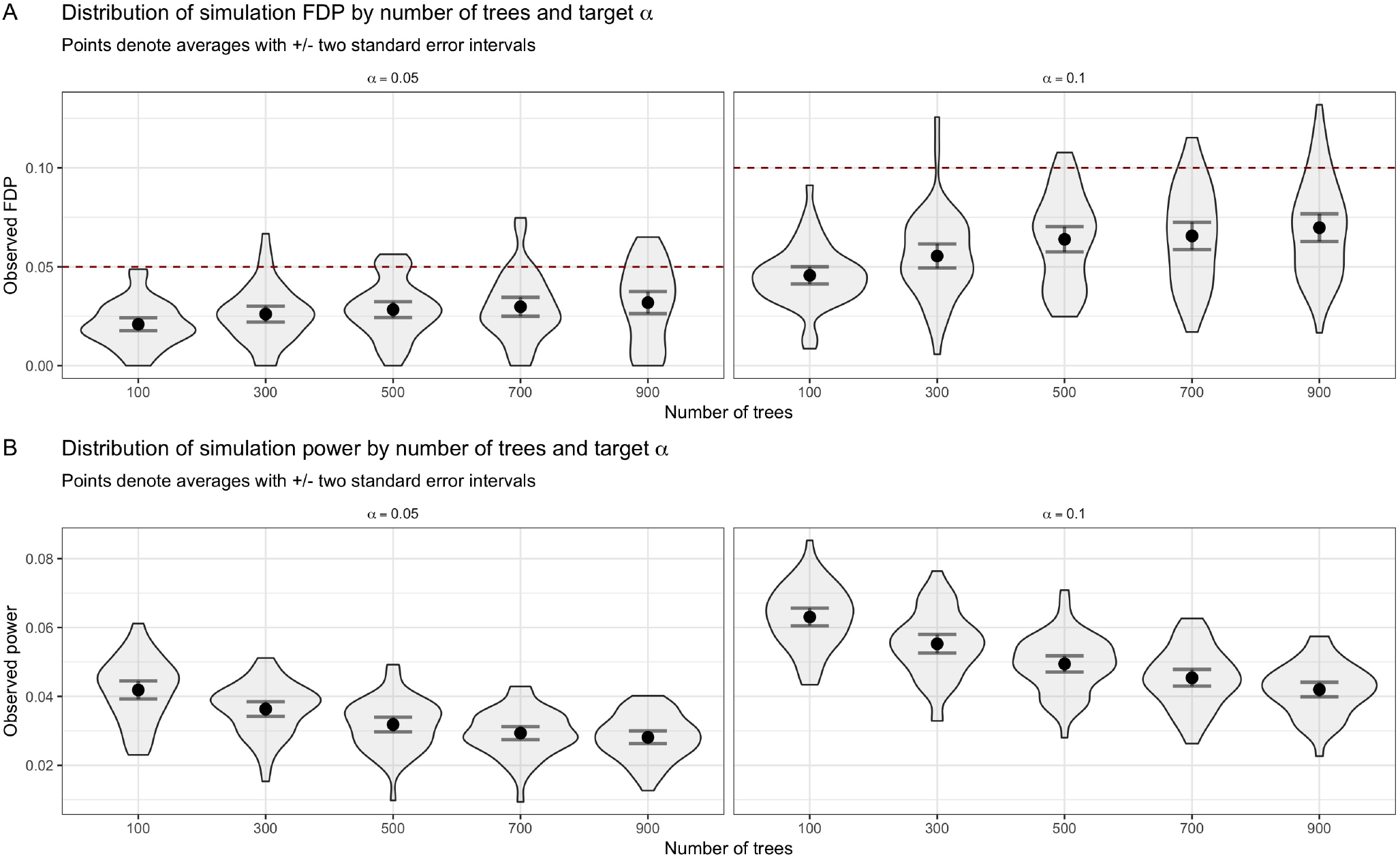
Distributions of observed *(A)* FDP and *(B)* power for simulations as the number of AdaPT gradient boosted trees increases by target FDR level *α*. Points denote averages with plus/minus two standard error intervals.

